# Widespread and targeted gene expression by systemic AAV vectors: Production, purification, and administration

**DOI:** 10.1101/246405

**Authors:** Rosemary C Challis, Sripriya Ravindra Kumar, Ken Y Chan, Collin Challis, Min J Jang, Pradeep S Rajendran, John D Tompkins, Kalyanam Shivkumar, Benjamin E Deverman, Viviana Gradinaru

## Abstract

We recently developed novel AAV capsids for efficient and noninvasive gene transfer across the central and peripheral nervous systems. In this protocol, we describe how to produce and systemically administer AAV-PHP viruses to label and/or genetically manipulate cells in the mouse nervous system and organs including the heart. The procedure comprises three separate stages: AAV production, intravenous delivery, and evaluation of transgene expression. The protocol spans eight days, excluding the time required to assess gene expression, and can be readily adopted by laboratories with standard molecular and cell culture capabilities. We provide guidelines for experimental design and choosing the capsid, cargo, and viral dose appropriate for the experimental aims. The procedures outlined here are adaptable to diverse biomedical applications, from anatomical and functional mapping to gene expression, silencing, and editing.

## INTRODUCTION

Recombinant adeno-associated viruses (AAVs) are commonly used vehicles for *in vivo* gene transfer and promising vectors for therapeutic applications^1^. However, AAVs that enable efficient and noninvasive gene delivery across defined cell populations are needed. Current gene delivery methods (e.g., intraparenchymal surgical injections) are invasive, and alternatives such as intravenous administration require high viral doses and still provide relatively inefficient transduction of target cells. We previously developed CREATE (Cre REcombination-based AAV Targeted Evolution) to engineer and screen for AAV capsids capable of more efficient gene transfer to specific cell types via the vasculature^2,3^. Compared to naturally occurring capsids, the novel AAV-PHP capsids exhibit markedly improved tropism for cells in the adult mouse central nervous system (CNS), peripheral nervous system (PNS), and visceral organs. In this protocol, we describe how to package genetic cargo into AAV-PHP capsids and intravenously administer AAVs for efficient and noninvasive gene delivery throughout the body (**Fig. 1**).

**Figure 1.**
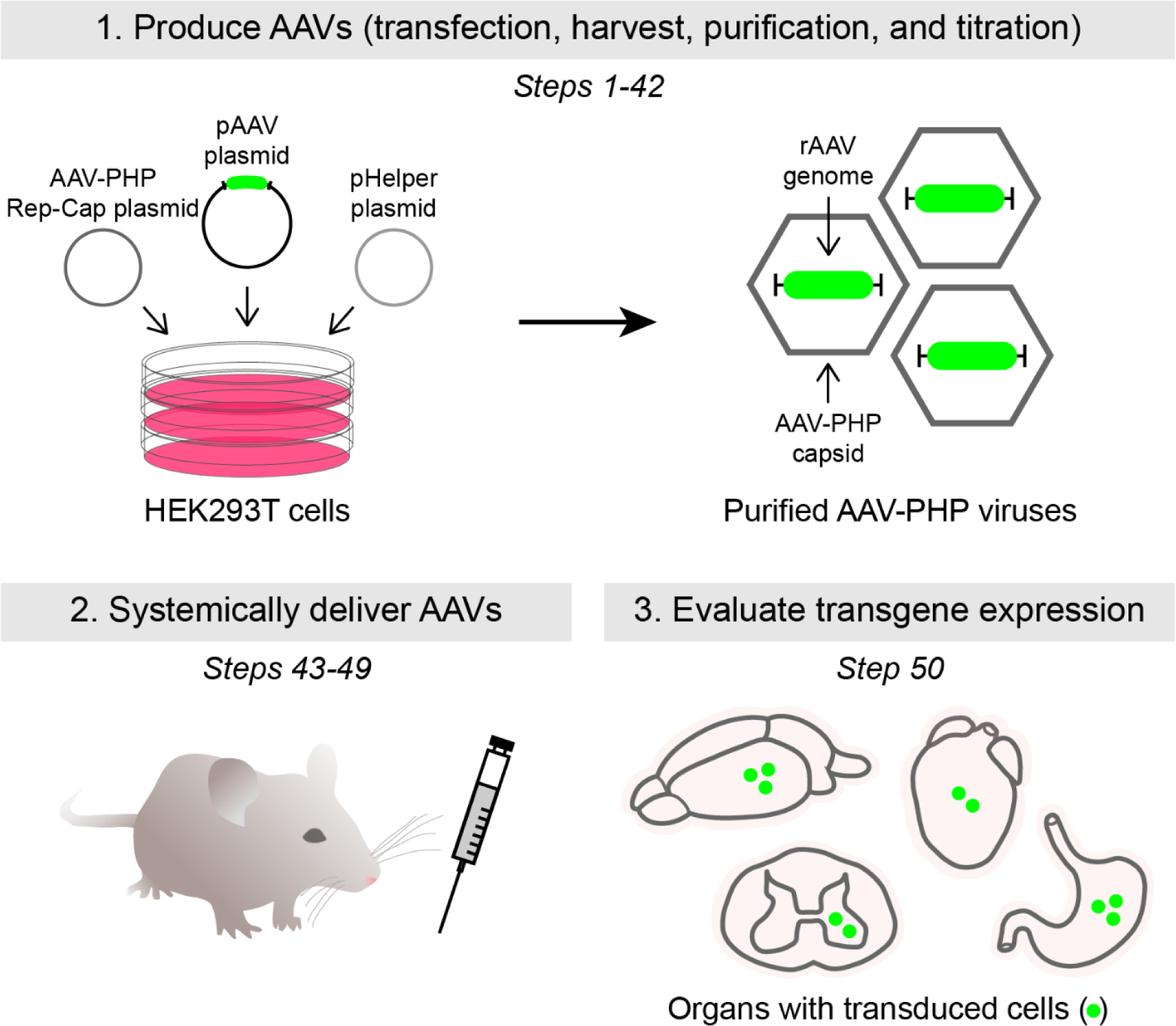
Overview of the protocol. The procedure comprises three main stages: AAV production (Steps 1-42), intravenous delivery (Steps 43-49), and evaluation of transgene expression (Step 50). The pAAV plasmid contains the rAAV genome (e.g., containing a fluorescent reporter, shown in green) (**Fig. 5** and **Table 1**), which is packaged into an AAV-PHP capsid via triple transient transfection. Systemic administration of AAV-PHP viruses is achieved via retro-orbital injection into wild-type or transgenic mice; transgene expression is evaluated after adequate time has passed for viral transduction and protein expression. AAV-PHP viruses target cells in the CNS (e.g., in the brain and spinal cord) or PNS and visceral organs (e.g., in the heart and gut). Filled green circles represent transduced cells. For illustrative purposes, we use fluorescent labeling as an example of how to assess transgene expression; however, assessment can take on other forms (see Experimental design section for details). See **Figure 6a** for a timeline of the procedure.

We recently identified several new capsid variants with distinct tropisms^2,3^. AAV-PHP.B and the further evolved AAV-PHP.eB efficiently transduce neurons and glia throughout the CNS (**Fig. 2**); a second variant, AAV-PHP.S, displays improved tropism for neurons within the PNS (**Fig. 3**) and organs including the gut^2^ and heart (**Fig. 4**). Importantly, these capsids target cell populations that are normally difficult to access due to their location (e.g., sympathetic, nodose, dorsal root, and cardiac ganglia) (**Fig. 3a-c** and **Fig. 4d**) or broad distribution (e.g., throughout the brain or enteric nervous system) (**Figs. 2** and **3d**). Together with the capsid, the genetic cargo (or AAV genome) can be customized to control transgene expression (**Fig. 5** and **Table 1**). The recombinant AAV (rAAV) genome contains the components required for gene expression including promoters, transgenes, protein trafficking signals, and recombinase-dependent expression schemes. Hence, different capsid-cargo combinations create a versatile AAV toolbox for genetic manipulation of diverse cell populations in wild-type and transgenic animals. Here, we provide researchers, especially those new to working with AAVs or systemic delivery, with resources to utilize AAV-PHP viruses in their own research.

**Figure 2.**
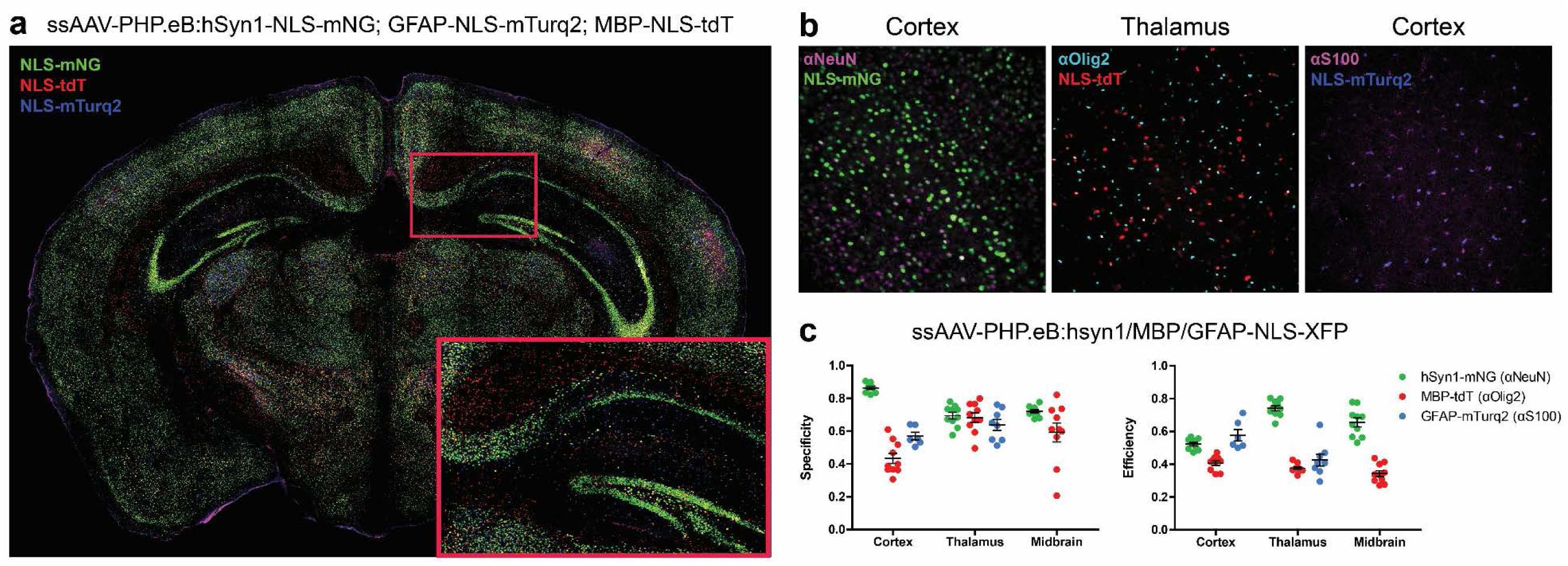
AAV-PHP.eB targets diverse cell types throughout the brain. We used AAV-PHP.eB to package single-stranded (ss) rAAV genomes that express nuclear localized (NLS) fluorescent reporters (XFP) from cell type-specific promoters. Genomes containing the hSyn1, MBP, or GFAP promoters were used to target neurons, oligodendrocytes, or astrocytes, respectively. Viruses were co-delivered via retro-orbital injection to a 7 week old wild-type mouse at 3 × 10^11^ vg/virus; gene expression in the brain was evaluated 4 weeks later. (**a**) Coronal image of the brain; inset shows zoomed-in view of the hippocampus. XFPs were mNeonGreen (mNG), tdTomato (tdT), or mTurquoise2 (mTurq2). (**b** and **c**) Antibody staining can be used to determine the specificity and efficiency of cell type-specific promoters. (**b**) NeuN, Olig2, and S100 mark neurons, oligodendrocyte lineage cells, and a population of glia that consists mostly of astrocytes, respectively. Images from the cortex and thalamus are shown. All brain slices were mounted in Prolong Diamond Antifade; images were acquired using confocal microscopy and are presented as maximum intensity projections. Refer to **Table 1** for details regarding rAAV genomes. (**c**) “Specificity” or “efficiency” are defined as the ratio of XFP labeled/antibody labeled cells to the total number of XFP or antibody labeled cells, respectively. For image processing, median filtering and background subtraction using morphological opening were first applied to each image to reduce noise and correct imbalanced illumination. Each nucleus expressing XFPs and labeled with antibodies was then segmented by applying a Laplacian of Gaussian filter to the pre-processed images. We considered cells that are both expressing XFPs and labeled with antibodies if the nearest center-to-center distance between blobs (nuclei or cell bodies) in two channels was less than 7 μm (half of the cell body size). Experiments on vertebrates conformed to all relevant governmental and institutional regulations and were approved by the Institutional Animal Care and Use Committee (IACUC) and the Office of Laboratory Animal Resources at the California Institute of Technology.

**Figure 3.**
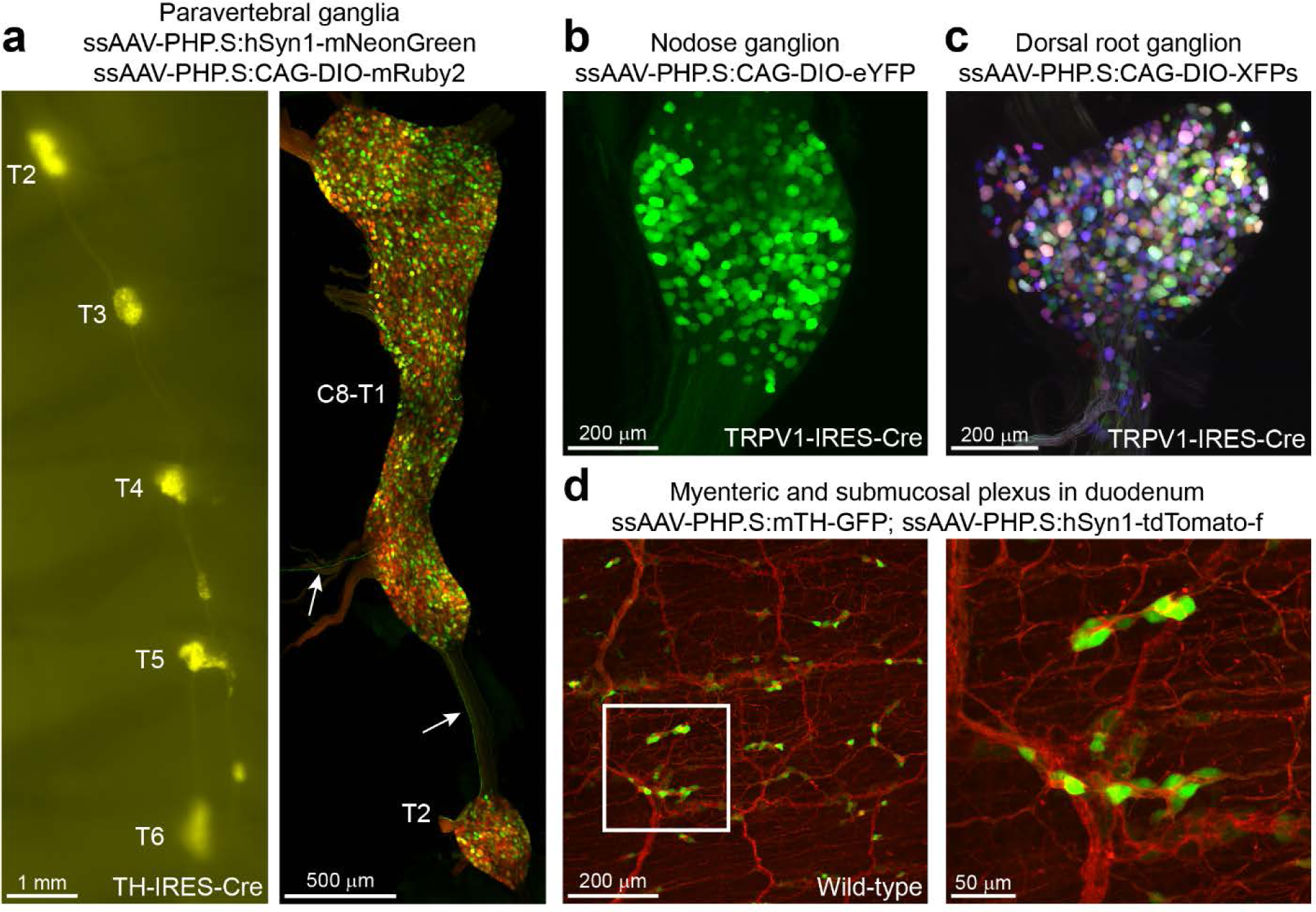
AAV-PHP.S transduces neurons throughout the PNS. We used AAV-PHP.S to package single-stranded (ss) rAAV genomes that express fluorescent reporters from either neuron-specific (e.g., hSyn1 and TH) or ubiquitous promoters (e.g., CAG). Viruses were delivered via retro-orbital injection to 6-8 week old wild-type or Cre transgenic mice and transgene expression was evaluated 2-3 weeks later. (**a**) ssAAV-PHP.S:hSyn1-mNeonGreen and ssAAV-PHP.S:CAG-DIO-mRuby2 were co-injected into a TH-IRES-Cre mouse at 1 × 10^12^ vg/virus. Native mNeonGreen and mRuby2 fluorescence were assessed 2 weeks later via wide-field (left) or confocal fluorescence microscopy (right). Images are from the second to sixth thoracic (T2-T6) (left) and eighth cervical to second thoracic (C8-T2) (right) paravertebral ganglia, which provide sympathetic innervation to thoracic organs including the heart. Arrows denote mNeonGreen-positive nerve fibers. (**b**) ssAAV-PHP.S:CAG-DIO-eYFP was injected into a TRPV1-IRES-Cre mouse at 1 × 10^12^ vg; gene expression in a nodose ganglion was evaluated 3 weeks later. (**c**) A mixture of 3 separate viruses (ssAAV-PHP.S:CAG-DIO-XFPs) were injected into a TRPV1-IRES-Cre mouse at 1 × 10^12^ vg/virus; gene expression in a dorsal root ganglion was evaluated 2 weeks later. XFPs were mTurquoise2, mNeonGreen, or mRuby2. (**d**) ssAAV-PHP.S:rTH-GFP and ssAAV-PHP.S:hSyn1-tdTomato-f (farnesylated) were co-injected into a C57BL/6J mouse at 5 × 10^11^ vg/virus; gene expression in the duodenum was assessed 22 d later. The image stack includes both the myenteric and submucosal plexus. Inset shows zoomed-in view of ganglia containing TH+ cell bodies; tdTomato-f labels both thick nerve bundles and individual fibers. Whole-mount tissues were optically cleared using either Sca/eSQ^25^ (**a** (right), **c**, and **d**) or RIMS^24^ (**b**) and imaged using confocal microscopy, unless stated otherwise; all confocal images are presented as maximum intensity projections. Refer to **Table 1** for details regarding rAAV genomes. Experiments on vertebrates conformed to allrelevant governmental and institutional regulations and were approved by the Institutional Animal Care and Use Committee (IACUC) and the Office of Laboratory Animal Resources at the California Institute of Technology.

**Figure 4.**
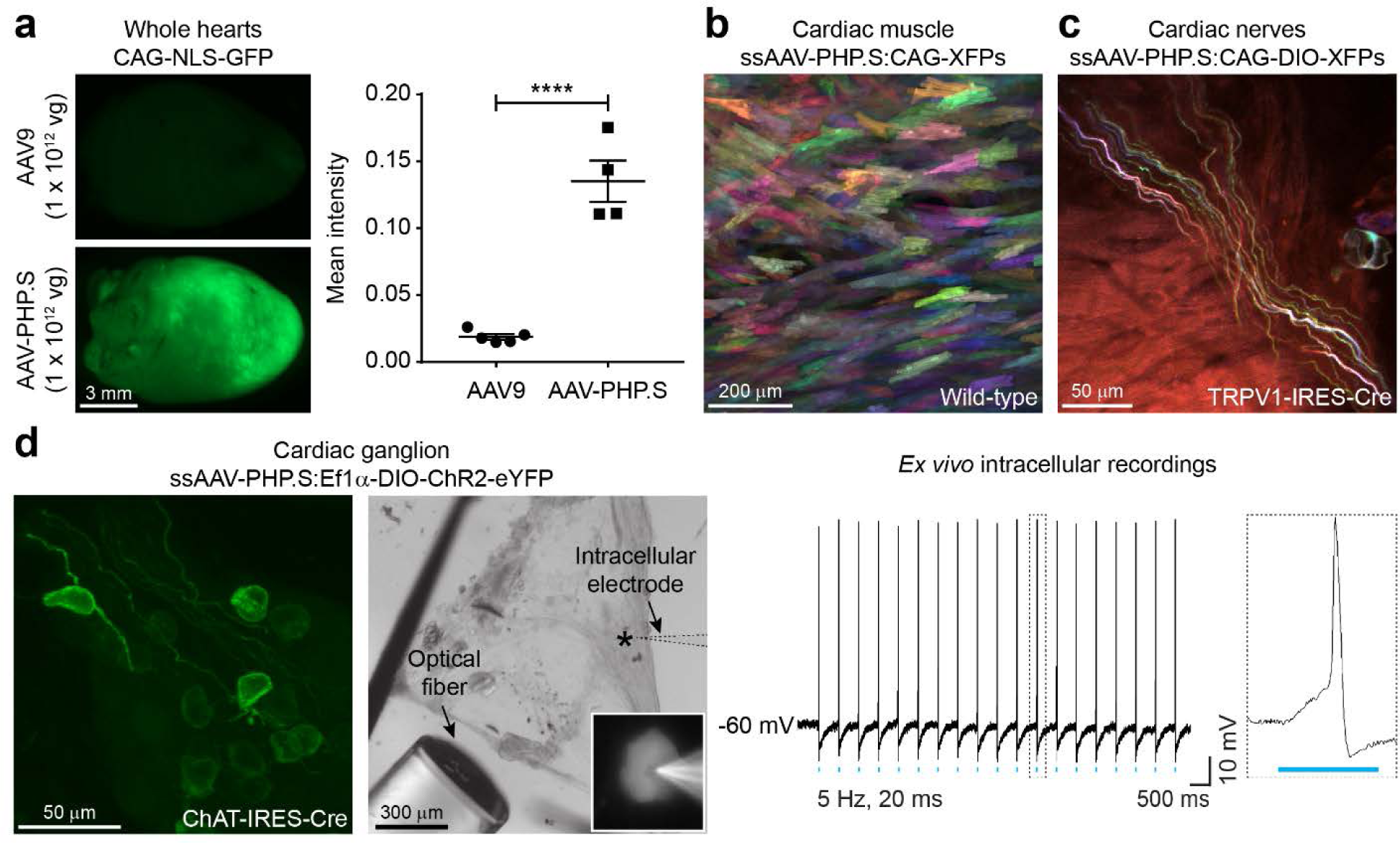
AAV-PHP.S for mapping the anatomy and physiology of the heart. AAV-PHP.S viruses were delivered via retro-orbital injection to 6-8 week old wild-type or Cre transgenic mice. (**a**) AAV-PHP.S transduces the heart more efficiently than the current standard, AAV9. ssAAV9:CAG-NLS-GFP or ssAAV-PHP.S:CAG-NLS-GFP were injected into C57BL/6J mice at 1 × 10^12^ vg. Native GFP fluorescence was assessed in whole mount hearts 4 weeks later via wide-field fluorescence microscopy (t_7_=8.449 and p<0.0001, unpaired *t* test). For AAV9 and AAV-PHP.S, n=5 and 4, respectively. (**b**) A mixture of 3 viruses (ssAAV-PHP.S:CAG-XFPs) were injected into a C57BL/6J mouse at 3.3 × 10^11^ vg/virus; gene expression in cardiac muscle was evaluated 11 d later, allowing individual cardiomyocytes to be easily distinguished from one another. XFPs were mTurquoise2, mNeonGreen, or mRuby2. (**c**) A mixture of 3 viruses (ssAAV-PHP.S:CAG-DIO-XFPs) were injected into a TRPV1-IRES-Cre mouse at 1 × 10^12^ vg/virus; gene expression in TRPV1-expressing cardiac nerves was evaluated 2 weeks later. (**d**) ssAAV-PHP.S:Ef1ɑ-DIO-ChR2-eYFP was injected into a ChAT-IRES-Cre mouse at 1 × 10^12^ vg; gene expression in a cardiac ganglion was evaluated 3 weeks later (left). *Ex vivo* intracellular recordings were performed after 5 weeks of expression. DIC image (middle) shows the optical fiber for light delivery and electrode for concurrent intracellular recordings; inset shows a higher magnification image of a selected cell (asterisk). Cholinergic neurons generated action potentials in response to 473 nm light pulses (5 Hz, 20 ms) (right). Whole-mount tissues were optically cleared using Sca/eSQ^25^ (**b**, **c**, and **d** (left)) and imaged using confocal microscopy, unless stated otherwise; all confocal images are presented as maximum intensity projections. Refer to **Table 1** for details regarding rAAV genomes. Experiments on vertebrates conformed to all relevant governmental and institutional regulations and were approved by the Institutional Animal Care and Use Committee (IACUC) and the Office of Laboratory Animal Resources at the California Institute of Technology.

**Figure 5.**
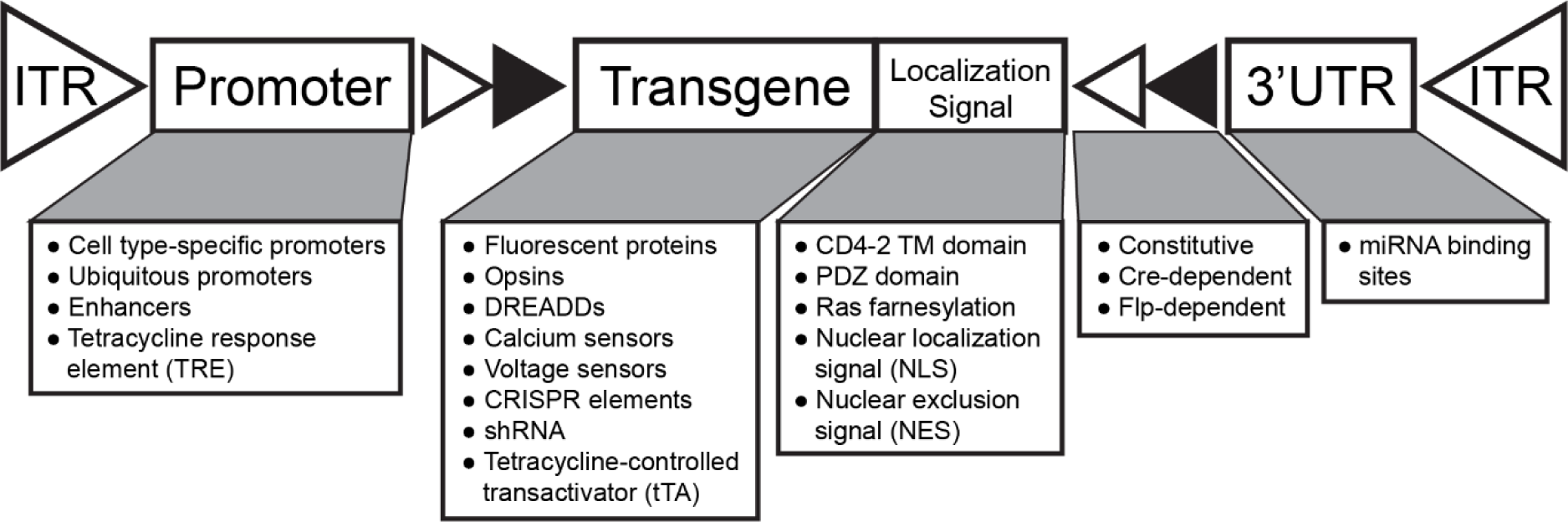
A modular AAV toolbox for cell type-specific gene expression. The rAAV genome, contained in a pAAV plasmid (not shown), consists of an expression cassette flanked by two 145 bp inverted terminal repeats (ITRs); the entire genome, including the ITRs, cannot exceed 5 kb. The promoters, transgenes, localization signals, and recombination schemes are interchangeable. Gene regulatory elements, such as promoters and microRNA (miRNA) binding sites, determine the strength and specificity of transgene expression16. Transgenes may be constitutively expressed or flanked by recombination sites for flippase (Flp)- or Cre recombinase (Cre)-dependent expression. In the latter approach, the transgene remains in the double-floxed inverted orientation (DIO); Cre- or Flp-mediated inversion of the transgene enables cell type-specific expression in transgenic mice (**Fig. 3a-c** and **4c-d**). Localization sequences further restrict gene expression to distinct cellular compartments such as the nucleus (via one or more nuclear localization signals (NLS)) (**Fig. 2**), cytosol (via a nuclear exclusion signal (NES)^58^), or cell membrane (via farnesylation55, the CD4-2^59^ transmembrane (TM) targeting domain, or PDZ^60^ protein-protein interaction domains) (**Fig. 3d**). Note that the 3’ UTR contains the woodchuck hepatitis posttranscriptional regulatory element (WPRE) (609 bp) and a polyadenylation signal (e.g., the human growth hormone (hGH) polyA) (479 bp) (not shown), both of which enhance transgene expression16. Use sequence editing and annotation software to determine the unique attributes of the rAAV genome. In **Table 1**, we list genomes used here and in our previous work^2,3^; see also Addgene’s plasmid repository for pAAVs that may be suitable for different applications. DREADDs = Designer Receptors Exclusively Activated by Designed Drugs; CRISPR = Clustered Regularly Interspaced Short Palindromic Repeats; shRNA = short hairpin RNA.

**Table 1.** AAV resources. A comprehensive list of plasmids used in this and related work^2,3^. The pHelper plasmid is available in Agilent’s AAV helper-free kit (Agilent, cat. no. 240071).

|  | Vector name pAAV- | Expression class | Addgene ID |
| --- | --- | --- | --- |
| Tunable expression | TRE-mTurq2 | tTA-dependent | 99113 |
|  | TRE-eYFP |  | 104056 |
|  | TRE-mRuby2 |  | 99114 |
|  | TRE-DIO-mTurq2 | Cre- and<br>tTA-dependent | 99115 |
|  | TRE-DIO-eYFP |  | 104057 |
|  | TRE-DIO-tdTomato |  | 99116 |
|  | TRE-DIO-mRuby2 |  | 99117 |
|  | CAG-tTA | Inducer | 99118 |
|  | hSyn1-tTA |  | 99119 |
|  | ihSyn1-tTA |  | 99120 |
|  | ihSyn1-DIO-tTA |  | 99121 |
| Tissue wide<br>expression | CAG-mTurq2 | Constitutive | 99122 |
|  | CAG-eYFP |  | 104055 |
|  | CAG-mRuby2 |  | 99123 |
|  | CAG-NLS-GFP-NLS |  | 104061 |
|  | CAG-DIO-mTurq2 | Cre-dependent | 104059 |
|  | CAG-DIO-eYFP |  | 104052 |
|  | CAG-DIO-mRuby2 |  | 104058 |
| Cell type-specific<br>expression | hSyn1-mTurq2 | Cell type-specific | 99125 |
|  | hSyn1-mRuby2 |  | 99126 |
|  | hSyn1-tdtomatoF |  | 104060 |
|  | MBP-NLS-tdTomato |  | 104054 |
|  | MsTH-GFP |  | 99128 |
|  | GFAP-NLS-mTurq2 |  | 104053 |
|  | GFAP-mKate2.5f |  | N/A |
|  | mDlx-NLS-mRuby2 |  | 99130 |

| **AAV-PHP Capsid** | ***In vivo* characteristics** | **Production** | **Addgene ID** |
| --- | --- | --- | --- |
| AAV-PHP.B | Broad CNS transduction | Good | 103002 |
| AAV-PHP.B2 | Broad CNS transduction | Good | 103003 |
| AAV-PHP.B3 | Broad CNS transduction | Good | 103004 |
| AAV-PHP.eB | Broad CNS transduction | Good | 103005 |
| AAV-PHP.S | Broad PNS transduction | Excellent | 103006 |
| AAV-PHP.A | Broad astrocyte transduction in CNS | Poor | N/A |

**Table 2.**
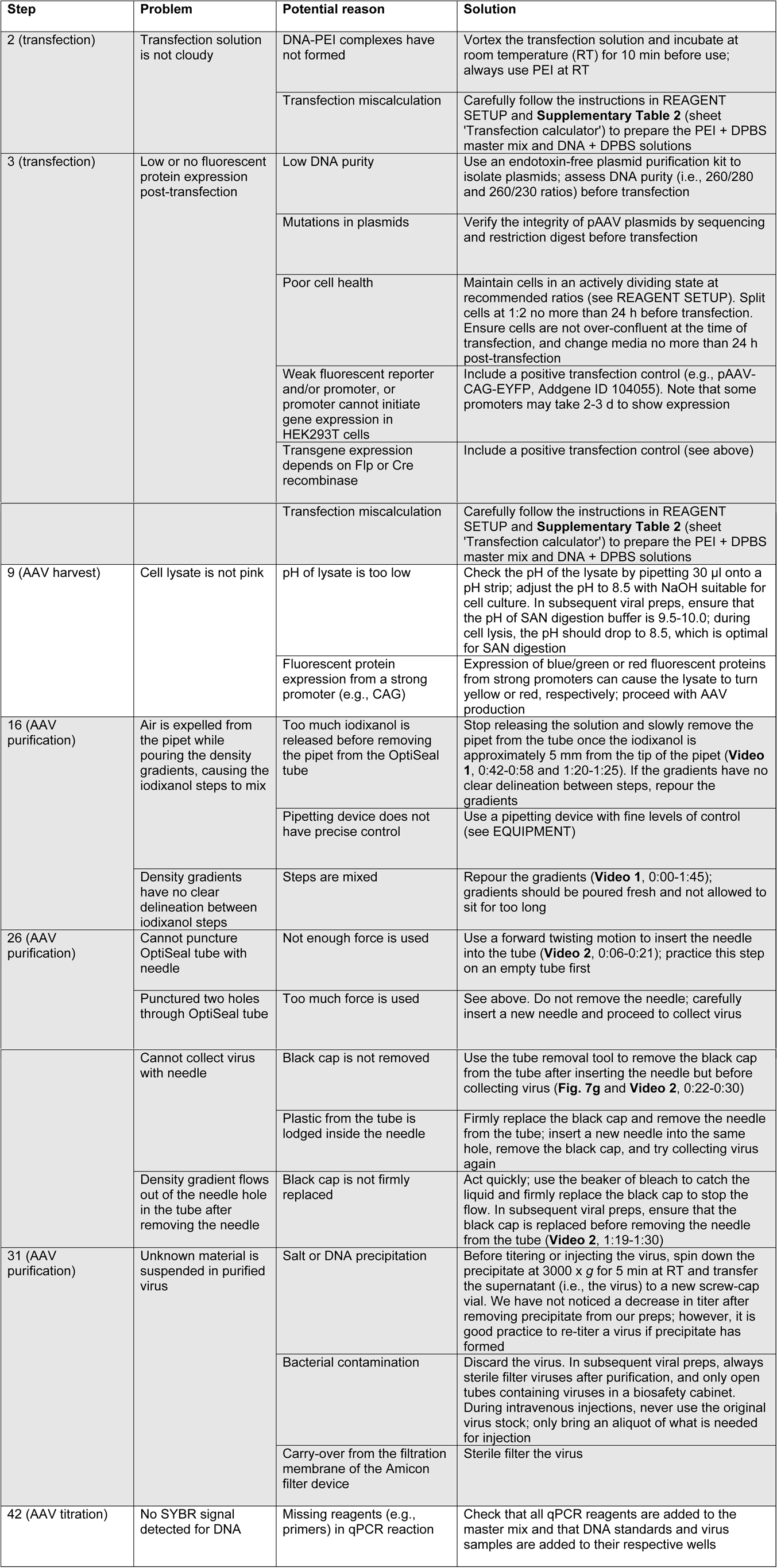
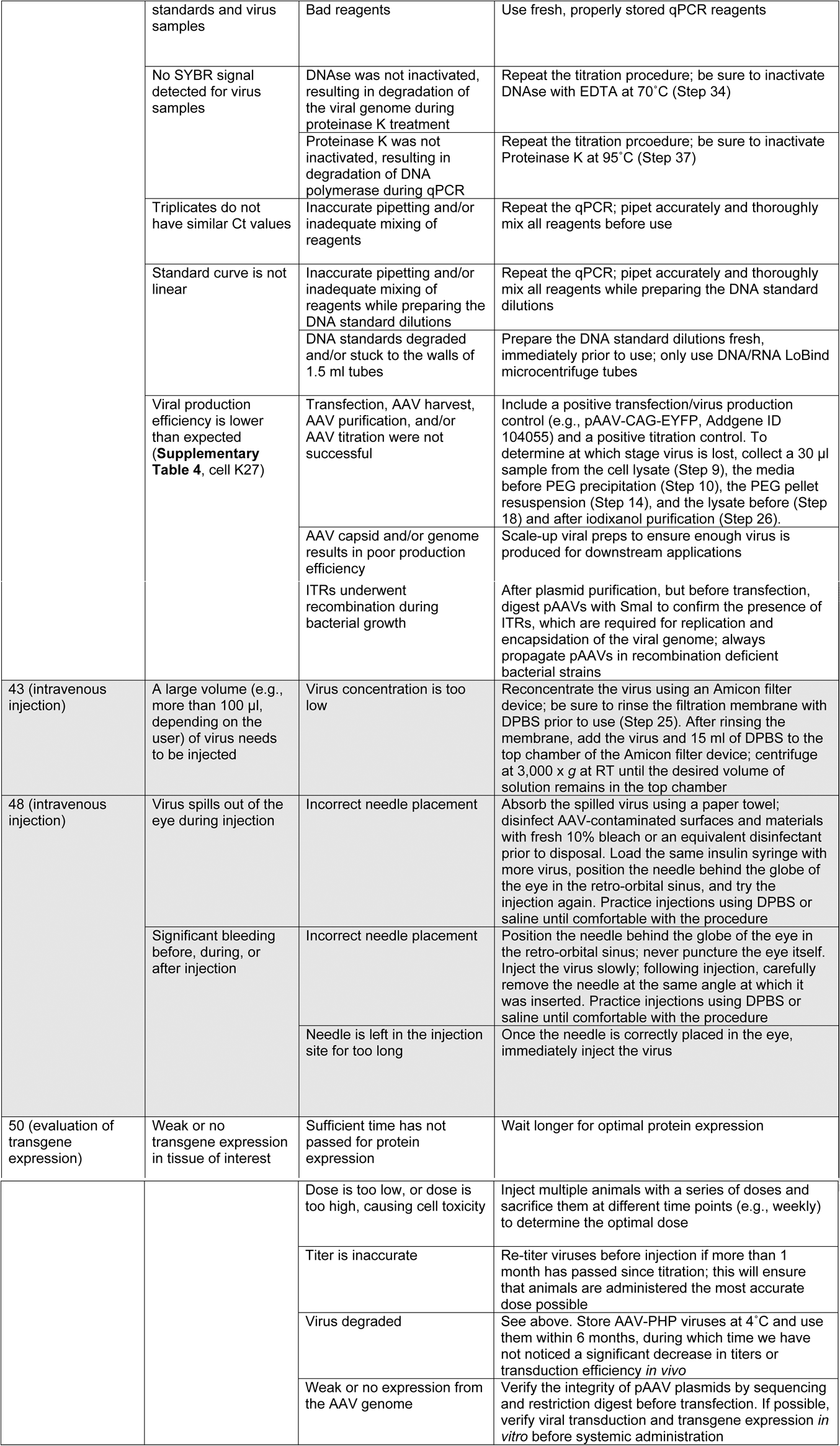
Troubleshooting AAV production, purification, and administration protocols.

### Overview of the protocol

We provide an instruction manual for users of AAV-PHP variants. The procedure includes three main stages (**Fig. 1**): AAV production (Steps 1-42), intravenous delivery (Steps 43-49), and evaluation of transgene expression (Step 50).

The AAV production protocol is adapted from established methods. First, HEK293T cells are transfected with three plasmids^4-6^ (Steps 1-3) (**Figs. 1** and **6**): (1) pAAV, which contains the rAAV genome of interest (**Fig. 5** and **Table 1**); (2) AAV-PHP Rep-Cap, which encodes the viral replication and capsid proteins; and (3) pHelper, which encodes adenoviral proteins necessary for replication. Using this triple transfection approach, the rAAV genome is packaged into an AAV-PHP capsid in HEK293T cells. AAV-PHP viruses are then harvested^7^ (Steps 4-14), purified^8,9^ (Steps 15-31), and titered^10^ (Steps 32-42) (**Fig. 6**). Purified viruses are intravenously delivered to mice via retro-orbital injection^11^ (Steps 43-49) and gene expression is later assessed using molecular, histological, or functional methods relevant to the experimental aims (Step 50).

**Figure 6.**
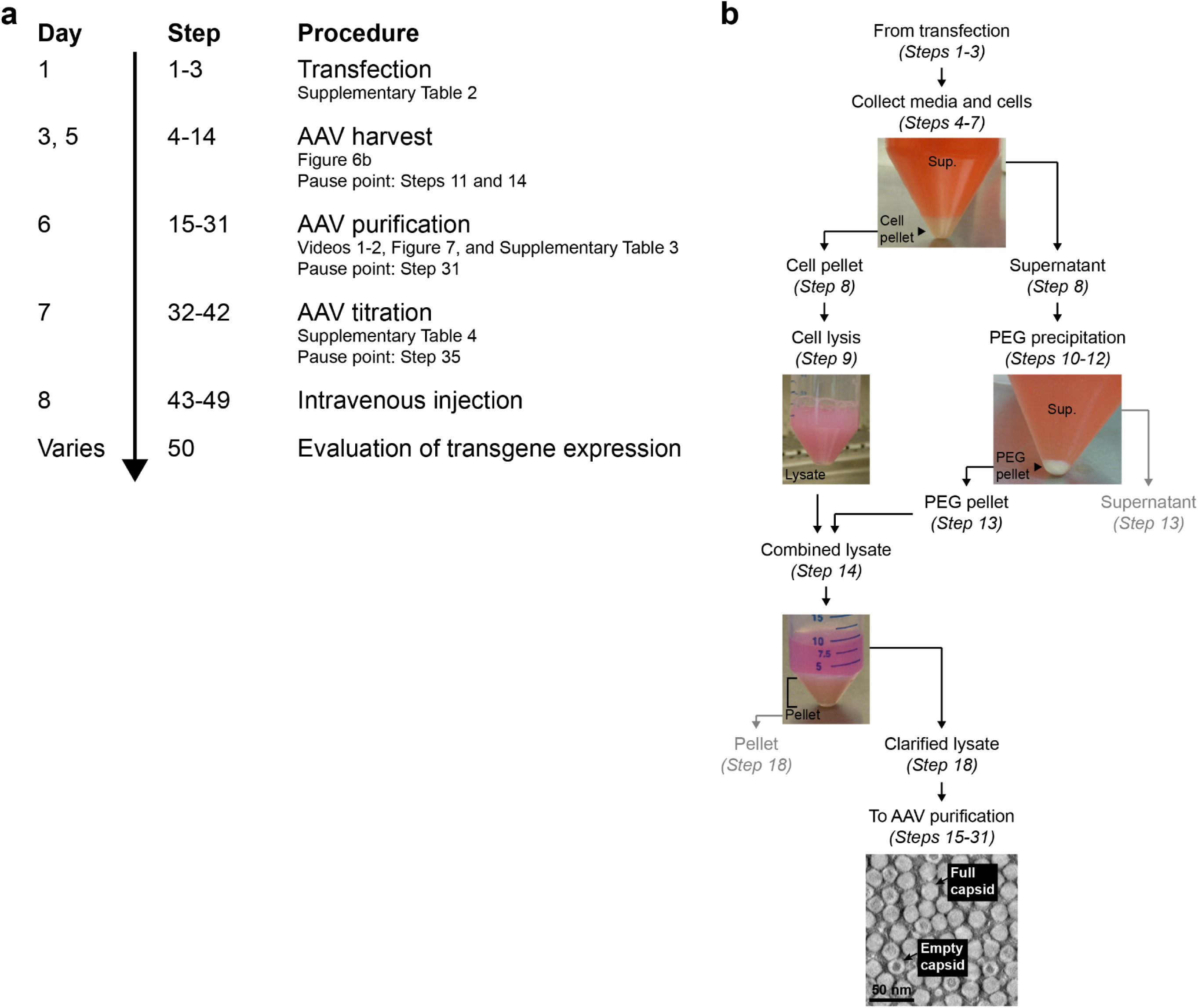
Timeline and AAV harvest procedure. **(a)** Timeline of the procedure. The entire protocol spans 8 d, excluding pause points on days 5 (Steps 11 and 14), 6 (Step 31), and 7 (Step 35) and the time required to evaluate transgene expression (Step 50). Days 1-7 (Steps 1-42) comprise the AAV production stage (**Fig. 1**). **(b)** Schematic of the AAV harvest procedure with images corresponding to indicated steps. Note that this iodixanol-based purification protocol does not eliminate empty capsids (i.e., capsids that fail to package a rAAV genome), as determined by negative staining transmission electron microscopy; empty particles are characterized by an electron-dense core. Grayed arrows and text denote steps where the supernatant and pellet can be bleached and discarded (Steps 13 and 18).

This protocol is optimized to produce AAVs at high titer (over 10^13^ vector genomes/ml) and with high transduction efficiency *in vivo*^2,3^.

### Experimental design

Before proceeding with the protocol, a number of factors should be considered, namely the expertise and resources available in the lab; the capsid and rAAV genome to be used; the dose for intravenous administration; and the method(s) available for assessing transgene expression. Each of these topics is discussed below and intended to guide users in designing their experiments.

#### Required expertise and resources

This protocol requires that scientists have basic molecular biology, cell culture, and animal work experience. Users should be approved to handle laboratory animals, human cell lines, and AAVs. A background in molecular cloning is advantageous though not necessary if relying on available plasmids.

In addition to having the above expertise, labs must be equipped for the molecular and cell culture work relevant to the procedure; we suggest that users read through the entire materials and procedure sections beforehand to ensure that the required reagents and equipment are available and appropriate safety practices and institutional approvals are in place.

#### Selecting an AAV-PHP capsid

We recommend choosing an AAV-PHP capsid based on its tropism and viral production efficiency. Capsid properties are listed in **Supplementary Table 1**; we include species, organs, and cell populations examined to date and note typical viral yields. We anticipate that most researchers will use AAV-PHP.eB (Addgene ID 103005) or AAV-PHP.S (Addgene ID 103006) in their experiments. AAV-PHP.eB and AAV-PHP.S produce viral yields similar to other high producing naturally occurring serotypes (e.g., AAV9) and enable efficient, noninvasive gene transfer to the CNS or PNS and visceral organs, respectively^2^ (**Figs. 2-4**).

The earlier capsid variants, which provide broad CNS transduction, either produce suboptimal yields (AAV-PHP.A)^3^ or have since been further evolved for enhanced transduction efficiency *in vivo* (AAV-PHP.B (Addgene ID 103002))^2^. We therefore recommend using AAV-PHP.eB for CNS applications, especially when targeting neurons. Note, however, that the chosen capsid will ultimately depend on the experimental circumstances; multiple factors including species^12^, age^13^, gender^14^, and health^15^ influence AAV tropism. Testing the AAV-PHP variants in a variety of experimental paradigms will continue to reveal the unique attributes of each capsid and identify those most suitable for different applications.

#### Selecting a rAAV genome

Users must select a rAAV genome, contained in a pAAV plasmid, to package into the capsid (**Figs. 1** and **5** and **Table 1**). In **Table 1**, we list pAAVs used here (**Figs. 2-4**) and in our previous work^2,3^; we direct users to Addgene’s plasmid repository for additional pAAVs developed for various applications.

Depending on the experimental aims, users may elect to design their own genomes^16^ and clone from existing pAAVs. When customizing plasmids, it is imperative that the rAAV genome, the sequence between and including the two inverted terminal repeats (ITRs), does not exceed 4.7-5 kb (**Fig. 5**); larger genomes will not be fully packaged into AAV capsids, resulting in truncated genomes and low titers. The ITRs are 145 base pair sequences that flank the expression cassette and are required for replication and encapsidation of the viral genome. ITRs are typically derived from the AAV2 genome and must match the serotype of the *rep* gene contained in the AAV Rep-Cap plasmid; AAV-PHP Rep-Cap plasmids contain the AAV2 *rep* gene and are therefore capable of packaging genomes with AAV2 ITRs. Other genetic components (e.g., promoters, transgenes, localization signals, and recombination schemes) are interchangeable and can be customized for specific applications (**Fig. 5**).

#### Dosage for intravenous administration

The optimal dose for intravenous administration to target cell populations must be determined empirically. We encourage users to consult **Figures 2-4** and related work for suggested AAV-PHP viral doses. The variants have been successfully employed for fluorescent labeling in adult mice^2,3,17^ (**Figs. 2-4**), neonatal mice^17^, and neonatal and adult rats^18^; they have also been administered for calcium imaging^19,20^ and optogenetic (**Fig. 4d**), chemogenetic^17^, and therapeutic applications^17,18^.

For applications using AAV-PHP.eB and AAV-PHP.S, we typically administer between 1 × 10^11^ and 3 × 10^12^ vector genomes (vg) of virus to adult mice (≥6 weeks of age). However, dosage will vary depending on the target cell population, desired fraction of transduced cells, and expression level per cell. AAVs independently and stochastically transduce cells, typically resulting in multiple genome copies per cell^2^. Therefore, higher doses generally result in strong expression (i.e., high copy number) in a large fraction of cells, whereas lower doses result in weaker expression (i.e., low copy number) in a smaller fraction of cells. To achieve high expression in a sparse subset of cells, users can employ a two-component system in which transgene expression is dependent on co-transduction of an inducer (e.g., a vector expressing the tetracycline-controlled transactivator (tTA))^2^; inducers are injected at a lower dose (typically 1 × 10^9^ to 1 × 10^11^ vg) to limit the fraction of cells with transgene expression. Note that gene regulatory elements (e.g., enhancers and promoters) also influence gene expression levels. Therefore, users should assess transgene expression from a series of doses and at several time points after intravenous delivery to determine the optimal experimental conditions.

#### Evaluation of transgene expression

Following *in vivo* delivery, AAV transduction and transgene expression increase over the course of several weeks. While expression is evident within days after transduction, it does not reach a steady state level until at least 3-4 weeks. Therefore, we suggest waiting a minimum of 2 weeks before evaluating fluorescent labeling^2,3,17,18^ (**Figs. 2-4**) and at least 3-4 weeks before beginning optogenetic (**Fig. 4d**), chemogenetic^17^, and calcium imaging^19,20^ experiments. Note that like other AAVs, AAV-PHP variants are capable of providing long-term transgene expression. AAV-PHP.B-mediated cortical expression of a genetically encoded calcium indicator, GCaMP6s, was reported to last at least 10 weeks post-injection without toxic side effects^20^ (i.e., nuclear filling^21^), and we have observed GFP expression throughout the brain more than one year after viral administration (see Supplementary Figure 4 in ref. ^3^). However, the time points suggested here are only meant to serve as guidelines; gene expression is contingent on multiple factors including the animal model, capsid, genome, and dose.

The appropriate method(s) for evaluating transgene expression will vary among users. Fluorescent protein expression can be assessed in thin or thick (≥100 μm) tissue samples. The CLARITY-based methods PACT (passive CLARITY technique) and PARS (perfusion-assisted agent release *in situ*)^22^ render thick tissues optically transparent while preserving their three-dimensional molecular and cellular architecture and facilitate deep imaging of large volumes (e.g., via confocal or light-sheet microscopy)^23^. Cleared tissues are compatible with endogenous fluorophores including commonly used markers like GFP^3,22,24^, eYFP22^22^, and tdTomato^24^. However, some fluorescent signals, such as those from mTurquoise2, mNeonGreen, and mRuby2, can deteriorate in chemical clearing reagents. To visualize these reporters, we suggest using optical clearing methods like RIMS (refractive index matching solution)^24^ or Sca/eSQ^25^ (**Figs. 3a and c**, and **4b-c**), or commercially available mounting media like Prolong Diamond Antifade (Thermo Fisher Scientific, cat. no. P36965)^2^ (**Fig. 2**). See the Anticipated Results section for details on expected outcomes when using fluorescent reporters.

We recognize that fluorescent labelling is not desired or feasible for every application. In such cases, users must identify the appropriate method(s) for examining transduced cells, which may include molecular (e.g., qPCR or Western blot), histological (e.g. with antibodies, small molecule dyes, or molecular probes) or functional (e.g., optical imaging) approaches.

### Limitations of the method

A major limitation of AAV capsids, including AAV-PHP variants, is their relatively small packaging capacity (<5 kb). Some elements of the rAAV genome, such as the WPRE (see legend in **Figure 5**), can be truncated^26^ or removed^27,28^ to accommodate larger genetic components. The development of smaller promoters^29,30^ and dual expression systems^31^, in which genetic elements are split between two or more viruses (requiring efficient cotransduction), have also enabled the delivery of larger genomes. Continued development of these approaches will help bypass restrictions on rAAV genome size.

Intravenous administration of AAVs also presents unique challenges. For example, systemic transduction may be undesirable for applications in which highly restricted gene expression is vital to the experimental outcome. Possible off-target transduction, due to the broad tropism of AAV-PHP variants and/or lack of compatible cell type-specific promoters, can be reduced by microRNA (miRNA)-mediated gene silencing. Sequences complementary to miRNAs expressed in off-target cell populations can be introduced into the 3’ UTR of the rAAV genome (**Fig. 5**); this has been shown to reduce off-target transgene expression and restrict expression to cell types of interest^32,33^.

Another challenge of systemic delivery is that it requires a high viral load, which can illicit an immune response against the capsid and/or transgene and reduce transduction efficiency *in vivo*^34^. Immunogenicity of AAVs may be exacerbated by empty capsid contamination in viral preparations^35,36^. The viral purification protocol (Steps 15-31) provided here reduces, but does not eliminate, empty capsids (**Fig. 6b**). If this poses a concern for specific applications, viruses can be purified using an alternative approach^7,8,37^.

Lastly, generating viruses for systemic administration may impose a financial burden on laboratories due to the doses of virus required. Nevertheless, viral-mediated gene delivery is inexpensive compared to creating and maintaining transgenic animals. Moreover, intravenous injection is faster, less invasive, and less technically demanding than other routes of AAV administration, such as stereotaxic injection, thereby eliminating the need for specialized equipment and survival surgery training.

### Applications of the method

We anticipate that AAV-PHP capsids can be used with the growing pAAV plasmid repository available through Addgene and elsewhere to enable a wide range of biomedical applications (**Fig. 5** and **Table 1**). Below, we highlight a few current and potential applications of this method.

#### Anatomical mapping

Fluorescent reporters are commonly used for cell type-specific mapping and phenotyping^2,38,39^ (**Figs. 2-4**). AAV-mediated multicolor labeling (e.g., Brainbow40) is especially advantageous for anatomical mapping approaches that require individual cells in the same population to be distinguished from one another. We and others have demonstrated the feasibility of this approach in the brain^2,40^, retina^40^, heart (**Figs. 4b and c**), and gut^2^, as well as peripheral ganglia (**Fig. 3c**). Spectrally distinct labeling is well-suited for studying the organization of cells (e.g., cardiomyocytes (**Fig. 4b**)) in healthy and diseased tissues and long-range tract tracing of individual fibers through extensive neural networks (e.g., the enteric^2^ or cardiac nervous systems (**Fig. 4c**)).

#### Functional mapping

AAV-PHP capsids are also relevant for probing cell function. AAV-PHP.B was previously used to target distinct neural circuits throughout the brain for chemogenetic^17^ and optical imaging applications^19,20^. We predict that AAV-PHP viruses will be beneficial for manipulating neural networks that are typically difficult to access, such as peripheral circuits controlling the heart (**Fig. 4d**), lungs^41^, or gut^42^. AAV-PHP variants could also be utilized to interrogate the function of non-neuronal cell types including cardiomyocytes^43^, pancreatic beta cells^44,45^, and hepatocytes^46^. Harnessing AAV-PHP viruses to modulate cell physiology may reveal novel roles for different cells in regulating organ function and/or animal behavior.

#### Gene expression, silencing, and editing

AAV-PHP viruses are well-suited for assessing potential therapeutic strategies that would benefit from organ-wide or systemic transgene expression. Recently, AAV-PHP.B was used to treat^17^ and model^18^ neurodegenerative diseases with widespread pathology. Other potential applications include gene editing (e.g., via CRISPR^47,48^) or silencing (e.g., via short hairpin RNA (shRNA)^49,50^); importantly, these approaches could be utilized to broadly and noninvasively manipulate cells in both healthy and diseased states for either basic research or therapeutically motivated studies.

### Summary

Systemically delivered AAV-PHP viruses provide efficient, noninvasive, and long-term gene transfer to cell types throughout the adult CNS, PNS, and visceral organs. Together with the use of custom rAAV genomes (**Fig. 5** and **Table 1**), researchers can target genetic elements to defined cell populations for diverse applications.

## MATERIALS

### REAGENTS

#### Plasmid DNA preparation

- Plasmids, supplied as bacterial stabs (Addgene; see **Table 1** for plasmids used in this and related work) CRITICAL Three plasmids (pAAV, capsid, and pHelper) are required for transfection (**Fig. 1**).
- Agarose (Amresco, cat. no. N605-250G)
- Antibiotics (e.g., carbenicillin disodium salt, Alfa Aesar, cat. no. J61949-06; all plasmids used in this work carry antibiotic resistance genes to ampicillin/carbenicillin)
- DNA ladder, 2-log, 0.1-10.0 kb (New England Biolabs, cat. no. N0550S)
- Lysogeny broth (LB) (Amresco, cat. no. J106-1KG); for large-scale plasmid preparations, such as maxi- and giga-preps, we typically use Plasmid+ media (Thomson Instrument Company, cat. no. 446300), an enriched media formulated to support higher cell densities and plasmid yields compared to LB
- Lysogeny broth (LB) with agar (Sigma-Aldrich, cat. no. L3147-1KG)
- NucleoBond Xtra Maxi Endotoxin-Free (EF) plasmid purification kit (Macherey-Nagel, cat. no. 740424.50) CRITICAL Triple transient transfection requires large amounts of capsid (22.8 μg/dish) and pHelper plasmid DNA (11.4 μg/dish) (**Supplementary Table 2**, sheet ‘Detailed calculations’); isolating these plasmids may be more convenient with a giga-scale purification kit (NucleoBond PC 10000 EF, Macherey-Nagel, cat. no. 740548). All plasmids should be purified under endotoxin-free conditions. Endotoxin contamination in plasmid preparations can reduce transfection efficiency, and contaminating endotoxins in viral preparations could elicit immune reactions in mammals *in vivo*.
- Restriction enzymes, including SmaI (New England Biolabs, cat. no. R0141S); used for verifying plasmid and ITR integrity
- Sequencing primers (Integrated DNA Technologies); used for verifying plasmid integrity
- SYBR Safe DNA gel stain (Invitrogen, cat. no. S33102)
- Tris-acetate-EDTA (TAE) buffer, 50X (Invitrogen, cat. no. B49)

#### Cell culture

- Human embryonic kidney (HEK) cells, 293 or 293T (ATCC, cat. no. CRL 1573 or CRL 3216) CAUTION HEK cells pose a moderate risk to laboratory workers and the surrounding environment and must be handled according to governmental and institutional regulations. Experiments involving HEK cells are performed using Biosafety Level 2 practices as required by the California Institute of Technology and the U.S. Centers for Disease Control and Prevention. The cell line identity has not been validated, nor do we routinely test for mycoplasma. CRITICAL HEK293 and HEK293T cells constitutively express two adenoviral genes, E1a and E1b, which are required for AAV production in these cells^6^; we do not recommend using an alternative producer cell line with this protocol.
- Dulbecco’s Modified Eagle Medium (DMEM), high glucose, GlutaMAX supplement, pyruvate (Gibco, cat. no. 10569-044)
- Ethanol, 70% (v/v); prepare from absolute ethanol (J.T. Baker, cat. no. 8025) CAUTION Ethanol is flammable.
- Fetal bovine serum (FBS) (GE Healthcare, cat. no. SH30070.03) CRITICAL Aliquot and store at −20°C for up to 1 year. Avoid freeze/thaw cycles.
- MEM Non-Essential Amino Acids (NEAA) solution, 100X (Gibco, cat. no. 11140-050)
- Penicillin-Streptomycin (Pen-Strep), 5000 U/ml (Gibco, cat. no. 15070-063) CRITICAL Aliquot and store at −20°C for up to 1 year. Avoid freeze/thaw cycles.
- TrypLE Express enzyme, 1X, phenol red (Gibco, cat. no. 12605-036)

#### Transfection

- Plasmid DNA CRITICAL We use a pAAV:capsid:pHelper plasmid ratio of 1:4:2 based on μg of DNA. We use 40 μg of total DNA per 150 mm dish (5.7 μg of pAAV, 22.8 μg of capsid, and 11.4 μg of pHelper) (**Supplementary Table 2**, sheet ‘Detailed calculations’).
- Polyethylenimine (PEI), linear, MW 25000 (Polysciences, Inc., cat. no. 23966-2)
- Water for Injection (WFI) for cell culture (Gibco, cat. no. A1287304)
- 1X Dulbecco’s PBS (DPBS), no calcium, no magnesium (Gibco, cat. no. 14190-250)
- 1 N HCl solution, suitable for cell culture (Sigma-Aldrich, cat. no. H9892) CAUTION HCl is corrosive. Use personal protective equipment.

#### AAV production

- Bleach, 10% (v/v); prepare fresh from concentrated liquid bleach (e.g., Clorox) CRITICAL AAV-contaminated materials must be disinfected with bleach prior to disposal; ethanol is not an effective disinfectant. Experiments involving AAVs follow a standard operating procedure in which contaminated equipment, surfaces, and labware are disinfected for 10 min with 10% bleach. AAV waste disposal should be conducted according to federal, state, and local regulations.
- Dry ice; optional
- KCl (any)
- MgCl_2_ (any)
- NaCl (any)
- OptiPrep (60% (w/v) iodixanol) density gradient media (Cosmo Bio USA, cat. no. AXS-1114542-5)
- Phenol red solution (Millipore, cat. no. 1072420100)
- Pluronic F-68 non-ionic surfactant, 100X (Gibco, cat. no. 24040-032); optional
- Polyethylene glycol (PEG), MW 8000 (Sigma-Aldrich, 89510-1KG-F)
- Salt-active nuclease (SAN) (ArcticZymes, cat. no. 70910-202)
- Tris base (any)
- Ultrapure DNAse/RNAse-free distilled water (Invitrogen, cat. no. 10977-023)
- Water for Injection (WFI) for cell culture (Gibco, cat. no. A1287304)
- 1X Dulbecco’s PBS (DPBS), no calcium, no magnesium (Gibco, cat. no. 14190-250)

#### AAV titration

- AAVs
- CaCl_2_ (any)
- DNAse I recombinant, RNAse-free (Sigma-Aldrich, cat. no. 4716728001)
- HCl, 37% (wt/wt) (Sigma-Aldrich, cat. no. 320331-500ML)
- MgCl_2_ (any)
- NaCl (any)
- *N*-lauroylsarcosine sodium salt (Sigma-Aldrich, cat. no. L9150-50G)
- Plasmid DNA containing the target sequence (e.g., pAAV-CAG-eYFP, Addgene ID 104055); used for preparing the DNA standard stock CRITICAL The plasmid used to make the DNA standard must contain the same target sequence as the pAAV plasmid used to generate virus. The target sequence must be within the rAAV genome; we typically amplify a portion of the WPRE in the 3’ UTR of pAAVs (see legend in **Fig. 5**).
- Primers corresponding to the target sequence (Integrated DNA Technologies)

> WPRE-forward: GGCTGTTGGGCACTGACAAT
>
> WPRE-reverse: CCGAAGGGACGTAGCAGAAG CRITICAL The proximity of the primer binding sites to the ITRs can affect titering results; therefore, titers measured with different primers or across laboratories may not be directly comparable.
- Proteinase K, recombinant, PCR grade (Sigma-Aldrich, cat. no. 03115828001)
- Qubit dsDNA HS assay kit (Invitrogen, cat. no. Q32854)
- ScaI-HF restriction enzyme (New England Biolabs, cat. no. R3122S) or other enzyme that cuts outside of the rAAV genome and within the pAAV backbone
- SYBR green master mix (Roche Diagnostics, cat. no. 04913850001)
- Tris base (any)
- Ultrapure DNAse/RNAse-free distilled water (Invitrogen, cat. no. 10977-023)
- Ultrapure EDTA, 0.5 M, pH 8.0 (Invitrogen, cat. no. 15575-020)

#### Intravenous (retro-orbital) injection

- AAVs
- Animals to be injected. This protocol describes the production of AAVs for intravenous delivery to 6-8 week old wild-type (C57BL/6J), ChAT-IRES-Cre (Jackson Laboratory, stock no. 028861, heterozygous), TH-IRES-Cre (European Mutant Mouse Archive, stock no. EM00254, heterozygous), and TRPV1-IRES-Cre mice (Jackson Laboratory, stock no. 017769, homozygous). CAUTION Experiments on vertebrates must conform to all relevant governmental and institutional regulations. Animal husbandry and experimental procedures involving mice were approved by the Institutional Animal Care and Use Committee (IACUC) and the Office of Laboratory Animal Resources at the California Institute of Technology.
- Bleach, 10% (v/v) prepared fresh, or equivalent disinfectant (e.g., Accel TB surface cleaner, Health Care Logistics, cat. no. 18692)
- Isoflurane, USP (Piramal Healthcare, 66794-017-25) CAUTION Isoflurane is a halogenated anesthetic gas associated with adverse health outcomes in humans and must be handled according to governmental and institutional regulations. To reduce the risk of occupational exposure during rodent anesthesia, waste gas is collected in a biosafety cabinet using a charcoal scavenging system as approved by the California Institute of Technology.
- Proparacaine hydrochloride ophthalmic solution, USP, 0.5% (Akorn Pharmaceuticals, cat. no. 17478-263-12)
- 1X Dulbecco’s PBS (DPBS), no calcium, no magnesium (Gibco, cat. no. 14190-250)

### EQUIPMENT

#### Plasmid DNA preparation equipment

- Autoclave (any, as available)
- Bunsen burner and lighter (Fisher Scientific, cat. nos. 03-917Q and S41878A)
- Centrifuge (see requirements for chosen plasmid purification kit)
- Gel electrophoresis system (Bio-Rad horizontal electrophoresis system)
- Gel imaging system (Bio-Rad Gel Doc EZ model)
- Incubating shaker (Eppendorf I24 model)
- Incubator (Thermo Fisher Scientific Heratherm model) or 37°C warm room
- Sequence editing and annotation software (e.g., Lasergene by DNASTAR, SnapGene by GSL Biotech, or VectorNTI by Thermo Fisher Scientific)
- Spectrophotometer (Thermo Fisher Scientific NanoDrop model)

#### Plasmid DNA preparation supplies

- Petri dishes, 100 mm × 15 mm (Corning, cat. no. 351029)
- Test tubes, 14 ml (Corning, cat. no. 352059)
- Ultra Yield flasks and AirOtop seals, 250 ml (Thomson Instrument Company, cat. nos. 931144 and 899423); use with Plasmid+ media. Alternatively, use LB and standard Erlenmeyer flasks.

#### AAV production equipment

- Biological safety cabinet CAUTION HEK293T cells and AAVs are biohazardous materials and must be handled according to governmental and institutional regulations. All experiments involving the aforementioned materials are performed in a Class II biosafety cabinet with annual certification as required by the California Institute of Technology and the U.S. Centers for Disease Control and Prevention.
- Centrifuge (any, provided the instrument can reach speeds up to 4,000 × *g*, refrigerate to 4°C, and accommodate 250 ml conical centrifuge tubes; we use the Beckman Coulter Allegra X-15R model)
- Fluorescence microscope for cell culture (Zeiss Axio Vert.A1 model)
- Incubator for cell culture, humidified at 37°C with 5% CO_2_ (Thermo Fisher Scientific Heracell 240i model)
- Laboratory balance (any, with a readability of 5-10 mg)
- Support stand with rod and clamp (VWR International, cat. nos. 12985-070, 60079-534, and 89202-624) (**Fig. 7f**)
- Ultracentrifuge (any preparative ultracentrifuge for *in vitro* diagnostic use; we use the Beckman Coulter Optima XE-9 model with the Type 70Ti rotor) CAUTION During ultracentrifugation, rotors are subjected to enormous forces (up to 350,000 × *g* in this protocol). Rotor failure can have catastrophic consequences including irreparable damage to the centrifuge and laboratory and fatal injuries to personnel. Inspect rotors for signs of damage or weakness prior to each use, and always follow the manufacturer’s instructions while operating an ultracentrifuge.
- Water bath (Fisher Scientific Isotemp model)

**Figure 7.**
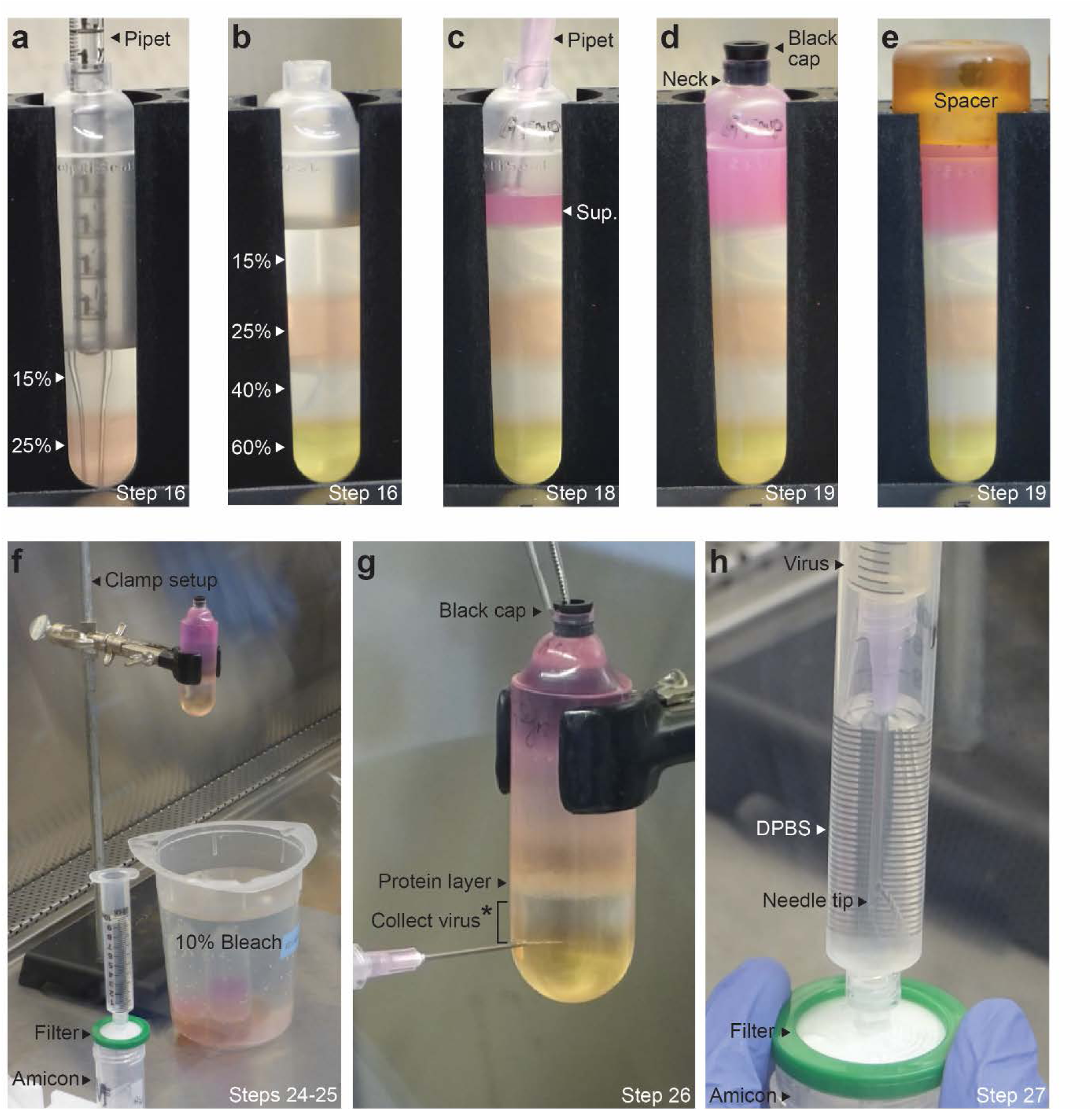
AAV purification procedure. (**a-b**) In Step 16, pipet the iodixanol density gradients (**Video 1**, 0:00-1:45). (**a**) Layer the 25% iodixanol underneath the 15% step (**Video 1**, 0:12-1:28). (**b**) Add layers of increasing density under the previous layer (**Video 1**, 1:29-1:45); the gradients should have a sharp delineation between steps. (**c**) In Step 18, load the supernatant (Sup.) from Step 17 (**Fig. 6b**) above the 15% step (**Video 1**, 1:46-2:22). (**d-e**) In Step 19, fill each tube up to the neck with SAN digestion buffer and insert a black cap (**d**); place a spacer on top before weighing the tubes (**e**). (**f**) After ultracentrifugation (Step 22), secure the tube into the clamp setup above a container of 10% bleach (Step 24). Allow 10 ml of DPBS to drip through the syringe filter unit into a rinsed Amicon filter device (Step 25). (**g**) In Step 26, collect the virus (**Video 2**, 0:00-1:30). Insert the needle approximately 4 mm below the 40/60% interface (i.e., where the tube just starts to curve) (**Video 2**, 0:06-0:21). Do not collect virus until the black cap is removed (asterisk) (**Video 2**, 0:22-0:30); do not collect from the protein layer at the 25/40% interface. (**h**) In Step 27, filter the virus/iodixanol (**Video 2**, 1:31-2:35). Inject the virus below the DPBS in the filter-attached syringe barrel (**Video 2**, 1:31-2:06) before pushing the virus/DPBS through the syringe filter unit and into the Amicon filter device.

#### AAV production supplies

- Amicon Ultra-15 centrifugal filter devices, 100 KDa molecular weight cutoff (Millipore, cat. no. UFC910024)
- Barrier pipet tips, 1000 μl (Genesee Scientific, cat. no. 23-430)
- Cell scrapers, 25 cm handle × 3 cm blade (Corning, cat. no. 353089)
- Conical centrifuge tubes, 50 ml and 250 ml (Corning, cat. nos. 352098 and 430776)
- Empty, sterile media bottles
- OptiSeal tubes (Beckman Coulter, cat. no. 361625); includes black caps
- OptiSeal tube kit (Beckman Coulter, cat. no. 361662); includes a tube rack, spacers, and spacer and tube removal tools
- Pipet Aid XL portable pipetting device (Drummond, cat. no. 4-000-105) CRITICAL Use a pipetting device with precise control, which is essential for pouring the density gradients in Step 16.
- pH indicator strips (Millipore, cat. nos. 109532 and 109584)
- Screw-cap vials, 1.6 ml (National Scientific Supply Co., cat. no. BC16NA-PS)
- Serological pipets, 2 ml, 5 ml, 10 ml, 25 ml, and 50 ml (Corning, cat. no. 356507 and Genesee Scientific, cat. nos. 12-102, 12-104, 12-106, and 12-107) CRITICAL Only Corning brand 2 ml serological pipets consistently fit into OptiSeal tubes while pouring the density gradients in Step 16. Alternatively, attach a small piece of clear tubing (6 mm) (e.g., Tygon Tubing) to a 5 ml pipet to pour the gradients.
- Stericup sterile vacuum filtration system, 0.22 μm, 500 ml and 1 L (Millipore, cat. nos. SCGPU05RE and SCGPU11RE)
- Sterile bottles, 500 ml (VWR International, cat. no. 89166-106)
- Syringes, 5 ml and 10 ml (BD, cat. nos. 309646 and 309604)
- Syringe filter unit, 0.22 μm (Millipore, cat. no. SLGP033RS)
- Tissue culture dishes, 150 mm × 25 mm (Corning, cat. no. 430599)
- 16 G × 1 ½ in needles (BD, cat. no. 305198)

#### AAV titration equipment

- Centrifuge (Eppendorf, 5418 model)
- Dry bath and heating blocks (Fisher Scientific Isotemp models)
- PCR plate spinner (VWR International, cat. no. 89184) or centrifuge equipped with plate adapters
- Quantitative PCR machine (any)
- Qubit 3.0 fluorometer (Invitrogen, cat. no. Q33216)

#### AAV titration supplies

- Barrier pipet tips, 10 μl, 20 μl, 200 μl, and 1000 μl (Genesee Scientific, cat. nos. 23-401, 23-404, 23-412, and 23-430)
- DNA clean-up kit, for purification of up to 25 μg of DNA standard (Zymo Research, cat. no. D4033)
- Microcentrifuge tubes, 1.5 ml DNA/RNA LoBind (Eppendorf, cat. no. 86-923)
- Qubit assay tubes (Invitrogen, cat. no. Q32856)
- Sealing film for 96-well PCR plates (Genesee Scientific, cat. no. 12-529)
- Stericup sterile vacuum filtration system, 0.22 μm, 250 ml (Millipore, cat. no. SCGPU02RE)
- Sterile bottles, 250 ml (VWR International, cat. no. 89166-104)
- 96-well PCR plates (Genesee Scientific, cat. no. 24-310W)

#### Intravenous (retro-orbital) injection equipment

- Animal anesthesia system (VetEquip, cat. nos. 901806, 901807, or 901810) CRITICAL Most animal facilities provide anesthesia systems equipped with an induction chamber, isoflurane vaporizer, nose cone, and waste gas scavenging system. In our experience, a mobile anesthesia system is most convenient for administering AAVs in a biosafety cabinet.

#### Intravenous (retro-orbital) injection supplies

- Activated charcoal adsorption filters (VetEquip, cat. no. 931401)
- Insulin syringes with permanently attached needles, 31 G × 5/16 in (BD, cat. no. 328438)
- Oxygen gas supply (any)
- Screw-cap vials, 1.6 ml (National Scientific Supply Co., cat. no. BC16NA-PS)

### REAGENT SETUP

CRITICAL All solutions should be prepared in a biosafety cabinet using sterile technique and endotoxin-free reagents and supplies. Glassware, stir bars, and pH meters are not endotoxin-free; autoclaving does not eliminate endotoxins. To prepare solutions, use pH indicator strips, dissolve reagents by heating and/or inverting to mix, and use demarcations on bottles to bring solutions up to the final volume. Reagents can be weighed outside of a biosafety cabinet since all solutions are filter sterilized before use.

#### Plasmid DNA

Grow bacterial stocks in LB or Plasmid+ media containing the appropriate selective antibiotic. Use a large-scale endotoxin-free plasmid purification kit to isolate plasmids; elute plasmid DNA with the supplied Tris-EDTA (TE) buffer. Measure DNA purity and concentration using a spectrophotometer and freeze at −20°C or −80°C for up to several years.

CRITICAL Always verify the integrity of purified plasmids by sequencing and restriction digest before proceeding with downstream applications. pAAV plasmids contain inverted terminal repeats (ITRs) (**Fig. 5**) that are prone to recombination in *E. coli*. pAAVs should be propagated in recombination deficient strains such as NEB Stable (New England Biolabs, cat. no. C3040H), Stbl3 (Invitrogen, cat. no. C737303), or SURE 2 competent cells (Agilent, cat. no. 200152) to prevent unwanted recombination. After purification, pAAVs should be digested with SmaI to confirm the presence of ITRs, which are required for replication and encapsidation of the viral genome; use sequence editing and annotation software to determine expected band sizes. Note that it is difficult to sequence through the secondary structure of ITRs^51^; avoid ITRs when designing sequencing primers.

#### DMEM + 5% (v/v) FBS

Add 25 ml of FBS, 5 ml of NEAA, and 5 ml of Pen-Strep to a 500 ml bottle of DMEM. Invert to mix and store at 4°C for up to several months. The resulting cell culture media should have a final concentration of 5% (v/v) FBS, 1X NEAA, and 50 U/ml Pen-Strep.

#### Cell culture

Thaw HEK293T cells according to the manufacturer’s recommendations. Passage cells using either TrypLE Express enzyme or a standard trypsinization protocol for adherent cultures^52^. Seed cells in 150 mm tissue culture dishes with a final volume of 20 ml of DMEM + 5% FBS per dish. Maintain in a cell culture incubator at 37°C with 5% CO_2_. CRITICAL We suggest a passage ratio of 1:3 (i.e., divide one dish of cells into three new dishes of cells every other day) when expanding cells for viral production; split cells at 1:2 (or 6 × 10^4^ cells/cm^2^) 24 hr before transfection. Always use sterile technique.

#### PEI stock solution

Pipet 50 ml of WFI water into a 50 ml conical centrifuge tube for later use. Add 323 mg of PEI to the remaining 950 ml bottle of WFI water and adjust the pH to 2-3 by adding 1 N HCl suitable for cell culture, keeping track of the volume of HCl added. Heat in a 37°C water bath for several hours (or overnight) and occasionally shake to mix. Once dissolved, add reserved WFI water to a total volume of 1 L. Filter sterilize, aliquot to 50 ml conical centrifuge tubes, and store at −20°C for up to 1 year. We routinely freeze/thaw our PEI aliquots. CRITICAL Both our PEI stock solution recipe and PEI calculations (**Supplementary Table 2**, sheet ‘Detailed calculations’) are based on ref. ^4^.

#### PEI + DPBS master mix

Thaw PEI in a 37°C water bath. Bring PEI to RT and vortex to mix. Add PEI and DPBS to a 50 ml conical centrifuge tube and vortex again to mix. Use **Supplementary Table 2** (sheet ‘Transfection calculator’) to calculate the volumes of PEI (cell I9) and DPBS (cell J9) needed. CRITICAL Prepare fresh the day of transfection.

#### DNA + DPBS

Bring plasmid DNA to RT and briefly vortex to mix. For each viral prep, add DNA and DPBS to a 50 ml conical centrifuge tube and use a P1000 pipet to mix. Use **Supplementary Table 2** (sheet ‘Transfection calculator’) to calculate the quantities of DNA (e.g., cells E9+E11+E13) and DPBS (e.g., cell F9) needed. CRITICAL Prepare fresh the day of transfection. Re-measure plasmid DNA concentrations immediately prior to use; multiple freeze/thaw cycles may cause DNA degradation.

#### SAN digestion buffer

Add 29.22 g of NaCl, 4.85 g of Tris base, and 952 mg of MgCl2 to a 1 L bottle of WFI water and shake to mix. Filter sterilize and store at RT for up to several months. The resulting SAN digestion buffer should have a final pH of 9.5-10.0 and a final concentration of 500 mM NaCl, 40 mM Tris base, and 10 mM MgCl_2_.

#### SAN + SAN digestion buffer

Add 100 U of SAN per ml of SAN digestion buffer; pipet to mix. CRITICAL Prepare fresh prior to use.

#### 40% (w/v) PEG stock solution

Decant approximately 500 ml of WFI water into a 500 ml sterile bottle for later use. Add 146.1 g of NaCl to the remaining 500 ml bottle of WFI water and shake/heat until dissolved. Once completely dissolved, add 400 g of PEG and heat at 37°C for several hours to overnight. Add reserved WFI water to a total volume of 1 L. Filter sterilize and store at RT for up to several months. The resulting stock solution should have a final concentration of 2.5 M NaCl and 40% (w/v) PEG. CRITICAL Prepare in advance. To expedite the procedure, heat the solution at 65°C until the PEG is dissolved. The solution will appear turbid but no flecks of PEG should remain; the mixture will become clear upon cooling. CRITICAL Pre-wet the entire filter surface with a minimal volume of water prior to adding the solution. This solution is extremely viscous and will take 1-2 h to filter.

#### DPBS + high salt

Add 29.22 g of NaCl, 93.2 mg of KCl, and 47.6 mg of MgCl_2_ to a 500 ml bottle of DPBS and shake to mix. Filter sterilize and store at RT for up to several months. The resulting buffer should have a final concentration of 1 M NaCl (in addition to the salt in the DPBS), 2.5 mM KCl, and 1 mM MgCl_2_.

#### DPBS + low salt

Add 2.92 g of NaCl, 93.2 mg of KCl, and 47.6 mg of MgCl_2_ to a 500 ml bottle of DPBS and shake to mix. Filter sterilize and store at RT for up to several months. The resulting buffer should have a final concentration of 100 mM NaCl (in addition to the salt in the DPBS), 2.5 mM KCl, and 1 mM MgCl_2_.

#### Iodixanol density step solutions (15%, 25%, 40%, and 60% (w/v) iodixanol)

For each step, add iodixanol, DPBS + high salt or DPBS + low salt, and phenol red (if applicable) to a 50 ml conical centrifuge tube. Briefly invert or vortex to mix. Use **Supplementary Table 3** to determine the volumes of each reagent needed. The 25% and 60% steps contain phenol red, which turns the solutions red and yellow, respectively, and facilitates clear demarcation of the gradient boundaries (**Fig. 7**). CRITICAL Prepare fresh the day of AAV purification. Alternatively, prepare up to 1 d in advance; store under sterile conditions at RT and protect from light. Do not pour the density gradients until Step 16.

#### 1 M Tris-Cl stock solution

Pipet 80 ml of Ultrapure or WFI water into a 250 ml sterile bottle. Add 12.11 g of Tris base and 7 ml of concentrated HCl and shake to mix; fine adjust the pH to 7.5 by adding more concentrated HCl. Add water to a total volume of 100 ml. Filter sterilize and store at RT for up to several months.

#### DNAse digestion buffer

Decant 250 ml of Ultrapure water into a 250 ml sterile bottle. Add 55.5 mg of CaCl_2_, 2.5 ml of 1 M Tris-Cl stock solution, and 238 mg of MgCl_2_ and shake to mix. Filter sterilize and store at RT for up to several months. The resulting buffer should have a final concentration of 2 mM CaCl_2_, 10 mM Tris-Cl, and 10 mM MgCl_2_.

#### DNAse I + DNAse digestion buffer

Add 50 U of DNAse I per ml of digestion buffer; pipet to mix. CRITICAL Prepare fresh prior to use.

#### Proteinase K solution

Decant 250 ml of Ultrapure water into a 250 ml sterile bottle. Add 14.61 g of NaCl and shake to mix. Add 2.5 g of *N*-lauroylsarcosine sodium salt to the mixture and gently swirl to mix; *N*-lauroylsarcosine sodium salt is a surfactant and will generate bubbles during vigorous mixing. Filter sterilize and store at RT for up to several months. The resulting solution should have a final concentration of 1 M NaCl and 1% (w/v) *N*-lauroylsarcosine sodium salt.

#### Proteinase K + proteinase K solution

Add 100 μg of proteinase K per ml of solution; pipet to mix. CRITICAL Prepare fresh prior to use.

#### DNA standard stock

Set up a single 50 μl restriction digest reaction; use 60-80 U (3-4 μl) of ScaI (or other suitable enzyme) to linearize 20 μg of the plasmid DNA containing the target sequence. Run a small amount of the reaction on an agarose gel to ensure complete digestion. Purify the reaction using two DNA clean-up columns. Measure the DNA concentration (ng/μl) using a spectrophotometer. Dilute to approximately 5-10 × 10^9^ single-stranded (ss) DNA molecules/μl and use the Qubit assay to verify the concentration (ng/μl). Divide into 20 μl aliquots and freeze at −20°C for up to 1 year. CRITICAL Prior to preparing the standard, use sequence editing and annotation software to confirm that the plasmid contains a single ScaI site in the ampicillin resistance gene. Refer to ref. ^10^ and use **Supplementary Table 4** (cell B13) to calculate the number of ssDNA molecules in a given plasmid. We typically use pAAV-CAG-mNeonGreen to prepare the standard; following restriction digest we dilute the linearized plasmid to 10 ng/μl, which corresponds to 6.6 × 10^9^ ssDNA molecules/μl.

#### DNA standard dilutions

Prepare three sets of 8 (1:10) serial dilutions of the DNA standard stock. For each set, begin by pipetting 5 μl of the standard into 45 μl of Ultrapure water (standard #8). Mix by pipetting and proceed with the 7 remaining dilutions (standard #7 to standard #1). The final concentrations of the standard dilutions should range from 5-10 × 10^8^ (standard #8) to 5-10 × 10^1^ (standard #1) ssDNA molecules/μl. CRITICAL Prepare fresh in DNA/RNA LoBind microcentrifuge tubes immediately prior to use; at low concentrations, the linearized DNA is prone to degradation and/or sticking to the walls of the tube^10^. One 20 μl aliquot of the DNA standard stock will provide enough DNA for preparing the dilutions and verifying the concentration via the Qubit assay prior to qPCR.

#### qPCR master mix

Prepare a qPCR master mix for the total number of reactions (i.e., wells) needed. One reaction requires 12.5 μl of SYBR green master mix, 9.5 μl of Ultrapure water, and 0.5 μl of each primer (from a 2.5 μM stock concentration), for a total of 23 μl/well. Briefly vortex to mix. CRITICAL Prepare fresh prior to use.

### EQUIPMENT SETUP

#### Clamp setup for AAV purification

Attach the rod to the support stand. Secure the clamp approximately 25-30 cm above the stand (**Fig. 7f**).

#### Anesthesia system

Place the induction chamber, nose cone, and waste gas scavenging system (e.g., activated charcoal adsorption filters) inside a biosafety cabinet. We recommend using a mobile anesthesia system in which the isoflurane vaporizer and oxygen supply remain outside of the cabinet workspace. Connect the associated tubing such that the input is from the vaporizer/oxygen supply and the output is to the charcoal scavenging device^53^.

## PROCEDURE

CRITICAL The entire procedure spans 8 d, excluding pause points and the time required to evaluate transgene expression (**Fig. 6a**). There are no pause points between days 1 and 5, until Step 11; once cells have been transfected, AAVs are harvested on days 3 and 5. Plan accordingly during this time window.

### Triple transient transfection of HEK293T cells

TIMING 1-2 h

CRITICAL For capsids that package well (i.e., AAV9, AAV-PHP.B, AAV-PHP.eB, and AAV-PHP.S), the PEI transfection protocol typically yields over 1 × 10^12^ vector genomes (vg) per 150 mm dish, as measured post-purification^2,3^. Before proceeding with the protocol, determine the number of dishes needed per viral prep and use **Supplementary Table 2** (sheet ‘Transfection calculator’) to calculate the quantities of PEI, DPBS, and plasmid DNA required for transfection. Skip to Step 43 if custom AAVs were obtained elsewhere.

1. 24 h before transfection, seed cells in 150 mm dishes to attain 80-90% confluency the next day^52^.
2. At the time of transfection, make the PEI + DPBS master mix and the DNA + DPBS solution for each viral prep (see REAGENT SETUP and **Supplementary Table 2**, sheet ‘Transfection calculator’). Using a serological pipet, add the required volume of the PEI + DPBS master mix (e.g., ‘Transfection calculator’ cell G9) to the DNA + DPBS solution (e.g., ‘Transfection calculator’ cells E9+E11+E13+F9); immediately vortex the resulting transfection solution to mix. Add 2 ml of the transfection solution dropwise to each dish and swirl to mix before returning to the cell culture incubator. CRITICAL STEP The transfection solution will appear slightly cloudy due to the formation of DNA-PEI complexes^4,5^. CRITICAL STEP Users may opt to run a positive transfection/virus production control (e.g. pAAV-CAG-eYFP, Addgene ID 104055); this is especially important if using an untested rAAV genome. TROUBLESHOOTING
3. Change the media 12-24 h post-transfection by aspirating the old media and replacing it with 20 ml of fresh, warmed DMEM + 5% FBS. CRITICAL STEP Do not allow cells to remain without media for more than a few min. To protect cells from unwanted stress, we aspirate media from 5 plates at a time and promptly replace it with new media. PEI is moderately cytotoxic^5^ and cell death up to 20% is common. Do not allow the media to go unchanged for more than 24 h post-transfection. Failure to change media in a timely manner will result in poor cell health and low titers. CRITICAL STEP Beginning 72 h post-transfection, examine cells under a fluorescence microscope to assess fluorescent protein expression, if applicable. Note that expression from the rAAV genome does not necessarily correlate with final viral yield and will depend on the reporter and/or promoter under investigation. TROUBLESHOOTING

### AAV harvest

TIMING 5 d

CAUTION While not infectious, recombinant AAVs are potent gene delivery vehicles and must be handled according to governmental and institutional regulations. The safety of packaged transgenes (e.g., oncogenic genes) should be carefully considered. Perform all procedures in a certified biosafety cabinet and clean AAV-contaminated equipment, surfaces, and labware with fresh 10% bleach prior to disposal.

CRITICAL Carefully label all tubes and replace gloves, pipets, and cell scrapers between viral preps to avoid cross-contamination. Refer to **Figure 6b** for a schematic of the AAV harvest procedure.

1. Harvest the cell culture media 72 h (3 d) post-transfection. Tilt each dish at a 30° angle and use a 25 ml serological pipet to collect the media. Store in an empty media bottle or sterile 500 ml bottle at 4°C until Step 6. Replace the media with 20 ml of fresh, warmed DMEM + 5% FBS. CAUTION Tilt dishes away from the front grill of the biosafety cabinet to prevent media from spilling out of the biosafety cabinet. CRITICAL STEP To avoid cross-contamination, harvest the media from one viral prep at a time. CRITICAL STEP For AAV-PHP production in HEK293T cells, the media at 72 h post-transfection contains approximately 2 × 10^11^ vg per dish, or approximately 20% of the expected viral yield. Failure to collect and change media at this time point will result in poor cell health and low titers. If time is limited, media and cells can be harvested at 72 h rather than 120 h (Step 5); however, total yields will be reduced by 30-50%.
2. Harvest the media and cells 120 h (5 d) post-transfection. Use a cell scraper to gently scrape the cells into the media. After scraping the first dish, prop it at a 30° angle using an empty 1.5 ml microcentrifuge tube rack for support. Scrape down the residual cells and media such that they are pooled together. Return the dish lid and scrape the next plate; prop dishes up against one another along the length of the biosafety cabinet until scraping is complete. Use a 25 ml serological pipet to collect the media and cells; transfer to a 250 ml conical centrifuge tube. CAUTION Scrape cells with a forward motion (i.e., away from the front grill of the biosafety cabinet) to prevent media and cells from splashing out of the biosafety cabinet. If a spill does occur at this or any other step, immediately cover with paper towels and carefully saturate the towels with 10% bleach. CRITICAL STEP To avoid cross-contamination, harvest the media and cells from one viral prep at a time.
3. Combine the media collected at 72 h post-transfection (Step 4) with the media and cells collected at 120 h post-transfection (Step 5). For smaller viral preps (1-5 dishes), proceed to option A. For larger preps (6-10 dishes), proceed to option B.

A. **Harvest from 1-5 dishes**

i. Pour the media collected in Step 4 into the corresponding 250 ml tube of media and cells collected in Step 5.
B. **Harvest from 6-10 dishes**

i. Pour the media collected in Step 4 into a second, empty 250 ml tube. CRITICAL STEP Save the bottle from Step 4 for Step 8.
4. Centrifuge the media and cells at 2,000 × *g* for 15 min at RT. Ensure that the tube caps are tightly secured. Centrifugation will result in the formation of a cell pellet (**Fig. 6b**).
5. Pour off the supernatant (i.e., the clarified media) into the corresponding bottle from Step 4. Allow excess media to drip back down onto the beveled edge of the 250 ml tube; remove using a P1000 pipet and combine with the supernatant. Store the supernatant at 4°C until Step 10. CRITICAL STEP Failure to remove excess media from the pellet will cause several ml of media to dilute the SAN digestion buffer in Step 9.
6. Resuspend the pellet for cell lysis. Prepare 5 ml of SAN + SAN digestion buffer per viral prep.

A. **Harvest from 1-5 dishes**

i. Use a 5 ml serological pipet to gently resuspend the pellet in 5 ml of SAN + SAN digestion buffer; pipet into a 50 ml tube to finish resuspending the pellet (**Fig. 6b**).
ii. Incubate in a 37°C water bath for 1 h and store at 4°C until Step 14.
B. **Harvest from 6-10 dishes**

i. Use a 10 ml serological pipet to partially resuspend the smaller pellet in 5 ml of SAN + SAN digestion buffer. Pipet into the second 250 ml tube containing the larger pellet and resuspend together; pipet into a 50 ml tube to finish resuspending the pellet (**Fig. 6b**).
ii. Incubate in a 37°C water bath for 1 h and store at 4°C until Step 14. CRITICAL STEP Be sure to collect the entire pellet, which sticks to the walls and beveled edges of 250 ml tubes. Save the 250 ml tubes for Step 10. CRITICAL STEP The high salt content of SAN digestion buffer lyses cells, which release viral particles and nucleic acids into solution. Initially, the cell lysate may be viscous and difficult to pipet; SAN will degrade nucleic acids and reduce the viscosity after incubation at 37°C. The pH of the lysate will decrease to 8-9 or lower during cell lysis but the lysate should appear pink rather than yellow/orange due to residual phenol red (**Fig. 6b**). Note that expression of fluorescent proteins from strong promoters (e.g., CAG) can alter the color of the lysate. CRITICAL STEP Collect a 30 μl sample from the cell lysate for troubleshooting. TROUBLESHOOTING
7. Retrieve the supernatant collected in Step 8.

A. **Harvest from 1-5 dishes**

i. Pour the supernatant from Step 8 into the corresponding 250 ml tube from Step 9.
B. **Harvest from 6-10 dishes**

i. Equally divide the supernatant from Step 8 into the two corresponding 250 ml tubes from Step 9. CRITICAL STEP Collect a 30 μl sample from the media for troubleshooting.
8. Use a 25 ml or 50 ml serological pipet to add 1/5 volume of 40% PEG stock solution to the supernatant (i.e., the supernatant should contain a final concentration of 8% PEG solution). Tighten the cap and invert 10 times to mix. Incubate on ice for 2 h. CRITICAL STEP For AAV production in HEK293T cells, the cell culture media contains a significant fraction of the expected yield^54^. Failure to PEG precipitate AAV particles in the media will result in lower viral yields^7^. PAUSE POINT The PEG/media mixture may be incubated at 4°C overnight.
9. Centrifuge the PEG/media mixture at 4,000 × *g* for 30 min at 4°C. Centrifugation will result in the formation of a PEG pellet (**Fig. 6b**).
10. Pour off the supernatant (i.e., the PEG-clarified media) into a used media collection bottle for bleaching. Allow excess media to drip back down onto the beveled edge of the 250 ml tube; aspirate to remove.
11. Resuspend the PEG pellet. Prepare 1 ml of SAN + SAN digestion buffer per viral prep.

A. **Harvest from 1-5 dishes**

i. Use a P1000 pipet to resuspend the PEG pellet in 1 ml of SAN + SAN digestion buffer; pipet into the corresponding lysate from Step 9 (**Fig. 6b**).
ii. Incubate in a 37°C water bath for an additional 30 min.
B. **Harvest from 6-10 dishes**

i. Use a P1000 pipet to partially resuspend one of the PEG pellets in 1 ml of SAN + SAN digestion buffer. Pipet into the second 250 ml tube containing the second pellet and resuspend together; pipet into the corresponding lysate from Step 9 (**Fig. 6b**).
ii. Incubate in a 37°C water bath for an additional 30 min. CRITICAL STEP Resuspending the PEG pellet is difficult and will take approximately 5 min per pellet. Be sure to collect the entire pellet, some of which sticks to the walls and beveled edges of 250 ml tubes. Do not use a serological pipet to resuspend the pellet, which can become lodged within the barrel of the pipet. CRITICAL STEP Collect a 30 μl sample from the PEG pellet resuspension for troubleshooting. PAUSE POINT Store the lysate at 4°C overnight. Alternatively, use a dry ice-ethanol bath to freeze the lysate; store at −20°C for up to 1 week.

### AAV purification

TIMING 1 d

CRITICAL One iodixanol density gradient is sufficient to purify virus from up to 10, 150 mm dishes. If more than 10 dishes per prep are used, divide the lysate into more than one gradient. The AAV purification steps are most easily learned by visualization; refer to **Figure 7** and **Videos 1-2** for details.

1. Determine the number of gradients needed and prepare the iodixanol density step solutions (see REAGENT SETUP and **Supplementary Table 3**). Set the OptiSeal tubes in the rack provided in the OptiSeal tube kit; alternatively, use the long edge of a 50 ml tube Styrofoam rack. CRITICAL STEP Check OptiSeal tubes for defects; do not use tubes with dents.
2. Pour the density gradients (**Fig. 7a-b** and **Video 1**, 0:00-1:45). Each gradient is composed of the following density steps: 6 ml of 15% iodixanol, 6 ml of 25% iodixanol, 5 ml of 40% iodixanol, and 5 ml of 60% iodixanol (**Supplementary Table 3**). Note that only Corning brand 2 ml serological pipets consistently fit into OptiSeal tubes; alternatively, attach a small piece of clear tubing (6 mm) (e.g., Tygon Tubing) to a 5 ml pipet to pour the gradients. Begin by pipetting 6 ml (measure to the 3 ml mark twice if using 2 ml pipets) of 15% iodixanol into each tube. Next, add 6 ml of 25% iodixanol under the 15% step by lightly touching the pipet tip to the bottom of the tube and slowly releasing the solution (**Fig. 7a**). Continue adding layers of increasing density under the previous layer. CRITICAL STEP Use a pipetting device with precise control. When loading the 25%, 40%, and 60% steps, stop releasing the solution and slowly remove the pipet once the iodixanol is approximately 5 mm from the tip of the pipet (**Video 1**, 0:42-0:58 and 1:20-1:25). This will prevent an air bubble from disturbing the gradient. The remaining iodixanol will be released when the pipet is removed from the tube. The gradients should have a sharp delineation between steps (**Fig. 7b**). TROUBLESHOOTING
3. Centrifuge the lysate from Step 14 at 2,000 × *g* for 10 min at RT. Centrifugation will result in the formation of a pellet (**Fig. 6b**).
4. Use a 2 ml serological pipet to load the supernatant (i.e., the clarified lysate) (approximately 6-7 ml total) from Step 17 above the 15% iodixanol step (**Fig. 7c** and **Video 1**, 1:46-2:22). Touch the pipet tip to the surface of the 15% step and slowly release the lysate such that a layer forms on top. CRITICAL STEP Do not allow the lysate to drip from the pipet tip onto the 15% step; this will cause the lysate to mix with the gradient. CRITICAL STEP The pellet may be soft, making it difficult to retrieve all of the supernatant. After loading 6-7 ml of lysate above the 15% step, spin the lysate for an additional 15 min at 3,000 × *g*; load the remaining supernatant onto the lysate layer. Discard the pellet. CRITICAL STEP Collect a 30 μl sample from the lysate for troubleshooting.
5. Fill each tube up to the neck with SAN digestion buffer. Insert a black cap (**Fig. 7d**) and place a spacer on top (**Fig. 7e**). Caps and spacers are provided with the OptiSeal tubes and in the OptiSeal tube kit, respectively.
6. Weigh the tubes with caps and spacers on. Balance the tubes to within 5-10 mg of each other using SAN digestion buffer. Be sure to adjust the tube weight in the biosafety cabinet; use the tube removal tool provided with the OptiSeal tube kit to remove the black cap and add the appropriate volume of SAN digestion buffer with a P20 or P200 pipet. CAUTION Failure to balance tubes before ultracentrifugation could result in damaged equipment.
7. Place the ultracentrifuge rotor in the biosafety cabinet. Load the tubes and fasten the lid. CAUTION To prevent damage to the rotor, set it on a paper towel so that the overspeed disk at the bottom is not scratched.
8. Carefully transfer the rotor to the ultracentrifuge. Spin the Type 70 Ti rotor at 350,000 × *g* (58,400 rpm) for 2 h and 25 min at 18°C with slow acceleration (#3) and deceleration (#9) profiles. Alternatively, a Type 60 Ti rotor can be used at 358,000 × *g* (59,000 rpm). CAUTION Always follow the manufacturer’s instructions while operating an ultracentrifuge.
9. During ultracentrifugation, gather the supplies and equipment for Steps 24-27. Assemble the clamp setup and collect one of each of the following per gradient: Amicon Ultra-15 centrifugal filter device, 5 ml syringe, 10 ml syringe, 0.22 μm syringe filter unit, and a 16 G needle.
10. After the spin is complete, bring the rotor inside the biosafety cabinet and remove the lid. Use the spacer removal tool provided with the OptiSeal tube kit to remove the spacer from the first tube. Next, use the tube removal tool to grip the tube neck. Slowly remove the tube from the rotor and secure it into the clamp setup above a 500 ml or 1 L beaker containing 10% bleach (**Fig. 7f**). Clean the side of the tube with 70% ethanol. CAUTION The black cap may become dislodged from the tube during removal, increasing the likelihood of a spill. Try replacing the cap before removing the tube from the rotor. Otherwise, replace the cap once the tube is secured in the clamp setup.
11. Prepare the supplies for Steps 26 and 27. First, rinse the filtration membrane in the Amicon centrifugal filter device by adding 15 ml of DPBS to the top chamber and centrifuging at 3,000 × *g* for 3 min at RT; discard the flow-through. Next, remove and save the plunger from a 10 ml syringe. Attach a 0.22 μm syringe filter unit to the syringe barrel and place it on top of a rinsed Amicon filter device. Add 10 ml of DPBS to the barrel and allow the solution to begin dripping through the syringe filter unit and into the filter device (**Fig. 7f**). Lastly, attach a 16 G needle to a 5 ml syringe.
12. Collect the virus from the 40/60% interface and 40% step^8,9^ (**Fig. 7g** and **Video 2**, 0:00-1:30). Hold the top of the OptiSeal tube with the non-dominant hand; use the dominant hand to hold the needle/syringe. Use a forward twisting motion to insert the needle approximately 4 mm below the 40/60% interface (i.e., where the tube just starts to curve). Use the tube removal tool in the non-dominant hand to remove the black cap from the tube to provide a hole for air entry. With the bevel up, use the needle/syringe to collect 4.0-4.5 ml of virus/iodixanol from the 40/60% interface and 40% step. Do not collect any of the visible protein band at the 25/40% interface; as this interface is approached, rotate the needle bevel down and continue collecting from the 40% step. Firmly replace the black cap before removing the needle from the tube. CAUTION Keep hands out of the path of the needle to prevent accidental exposure to AAVs. Failure to firmly replace the black cap before removing the needle will cause the AAV-contaminated solution to flow out of the needle hole in the tube and potentially onto and out of the biosafety cabinet. Perform this step over a large beaker of 10% bleach (**Fig. 7f**). CRITICAL STEP The virus should concentrate at the 40/60% interface and within the 40% step^9^. There will not be a visible virus band, but the phenol red in the 25% and 60% steps helps to better define the 40% cushion. CRITICAL STEP Practice puncturing an empty OptiSeal tube with a needle before attempting to collect virus from the density gradient. CRITICAL STEP Collect a 30 μl sample from the virus/iodixanol for troubleshooting. TROUBLESHOOTING
13. Add the 4.0-4.5 ml of virus/iodixanol to the syringe barrel containing 10 ml of DPBS (prepared in Step 25) (**Fig. 7h** and **Video 2**, 1:31-2:06). Layer the virus below the DPBS by placing the needle near the bottom of the barrel and pressing on the plunger. Insert the 10 ml syringe plunger into the barrel and push the virus/DPBS mixture through the syringe filter unit and into the Amicon filter device (**Video 2**, 2:07-2:35). Mix well using a P1000 pipet. CRITICAL STEP This filtration step reduces the likelihood of clogging the filtration membrane in the Amicon filter device. The virus/iodixanol mixture will be difficult to push through the syringe filter unit; DPBS will be easy to push through as it washes the filter. CRITICAL STEP AAVs adhere to hydrophobic surfaces including plastics. Pluronic F-68 is a non-ionic surfactant that may reduce virus loss associated with sticking to plastics. Optional: Include 0.001% Pluronic F-68 in DPBS for Steps 27-30.
14. Centrifuge the virus/DPBS mixture at 3,000 × *g* for 5-8 min, or until the volume of the solution remaining in the top chamber of the Amicon filter device is 500-1500 μl (>10X concentrated). CRITICAL STEP This step may take longer because iodixanol initially slows down the passage of the solution through the filtration membrane.
15. Discard the flow-through for bleaching. Add 14 ml of DPBS to the virus in the top chamber and use a P1000 pipet to mix.
16. Centrifuge the virus/DPBS mixture as in Step 28. Wash the virus 2 more times for a total of 4 buffer exchanges. During the last spin, retain 500-1000 μl of solution in the top chamber. CRITICAL STEP The third and fourth washes may only require a 2-3 min spin until the volume of the solution remaining in the top chamber is 500-1000 μl.
17. Filter sterilize the virus. Attach a 0.22 μm syringe filter unit to a 5 ml syringe. Pre-wet the filter with 200 μl of DPBS and discard the flow-through before filtering the virus. Use a P200 pipet to transfer the virus from the top chamber of the Amicon filter device directly into the syringe barrel. Push the virus through the filter and into a 1.6 ml screw-cap vial; store at 4°C. PAUSE POINT Store the purified virus under sterile conditions at 4°C for up to 6 months. CRITICAL STEP Amicon filter devices are not sterile; the virus should therefore be filter sterilized before storage. Virus stocks may be difficult to push through the filter; press the filter down onto the vial and push the plunger slowly to prevent liquid from spraying out of the syringe and onto or out of the biosafety cabinet. CRITICAL STEP The screw-cap vials are not low protein binding; however, they help prevent the formation of aerosols when opening and closing the tubes. We store AAVs in screw-cap vials at 4°C and typically use them within 3 months, during which time we have not noticed a significant decrease in titers or transduction efficiency *in vivo*. We have not rigorously tested the effects of long-term storage at −20°C or −80°C. TROUBLESHOOTING

### AAV titration

TIMING 1 d

CRITICAL The AAV titration procedure below is adapted from ref. ^10^. Each virus sample is prepared in triplicate in separate 1.5 ml DNA/RNA LoBind microcentrifuge tubes and later loaded into a 96-well plate for qPCR. All solutions must be accurately pipetted and thoroughly mixed; qPCR is highly sensitive to small changes in DNA concentration.

1. Prepare a plan for the PCR plate setup. Allocate the first 24 wells (A1-B12) for the DNA standards such that standard #1 occupies wells A1-A3, standard #2 occupies wells A4-A6, and so on. Use the remaining wells for the virus samples such that the first virus sample occupies wells C1-C3, the second sample occupies wells C4-C6, and so on. CRITICAL Include water as a negative control and a virus sample with a known concentration as a positive control; prepare the positive control with the virus samples in Steps 33-40.
2. Use DNAse I to digest DNA that was not packaged into the viral capsid. Prepare DNAse I + DNAse digestion buffer and aliquot 100 μl to each 1.5 ml tube. Vortex each virus for 1-2 s to mix and pipet 2 μl into each of three tubes. Briefly vortex and spin down; incubate in a 37°C water bath for 1 h. CRITICAL Prepare each virus sample in triplicate.
3. Inactivate the DNAse. Add 5 μl of EDTA to each tube; briefly vortex to mix, spin down, and incubate in a 70°C dry bath for 10 min. CRITICAL STEP DNAse must be inactivated or else it will degrade the viral genome when it is released from the viral capsid in Step 35.
4. Use proteinase K to digest the viral capsid and release the viral genome. Prepare proteinase K + proteinase K solution and add 120 μl to each tube. Briefly vortex and spin down; incubate in a 50°C dry bath for 2 h. PAUSE POINT Samples may be incubated at 50°C overnight.
5. During the last 20 min of the proteinase K digestion, prepare the DNA standard dilutions and use the Qubit assay to measure the concentration (ng/μl) of the DNA standard stock. CRITICAL STEP The concentration of the standard stock solution will be used to generate the standard curve after qPCR (**Supplementary Table 4**, cell B9).
6. Inactivate the proteinase K. Incubate the tubes in a 95°C dry bath for 10 min. CAUTION Tube caps may pop open unexpectedly; use safety glasses while removing the tubes from the 95°C dry bath. CRITICAL STEP Proteinase K must be inactivated or else it will digest DNA polymerase during qPCR.
7. Allow the tubes to cool. Briefly vortex each sample and add 3 μl to a new tube containing 897 μl of Ultrapure water (a 1:300 dilution). Briefly vortex the diluted samples.
8. Prepare the qPCR master mix.
9. Load the PCR plate based on the experimental plan from Step 32. First, pipet 23 μl of qPCR master mix into each designated well. Next, pipet 2 μl of each standard into wells A1-B12. Lastly, pipet 2 μl of each diluted sample from Step 38 into wells C1 and onward. Seal the plate with sealing film and briefly spin down in a plate spinner.
10. Place the PCR plate into the qPCR machine. Use the following cycling parameters: Step 1. 95°C 10 min Step 2. 95°C 15 s Step 3. 60°C 20 s Step 4. 60°C 40 s Repeat Steps 2-4 40X.
11. When the qPCR run is complete, export the cycle threshold (Ct) values to an Excel file. Use **Supplementary Table 4** to generate a standard curve and calculate the titer (vg/ml) (cell G27) of each virus; calculate per plate production (vg/dish) (cell K27) to assess production efficiency. TROUBLESHOOTING

### Intravenous (retro-orbital) injection

TIMING 2-5 min per mouse, excluding set-up and clean-up time

CAUTION Follow appropriate institutional and governmental guidelines and regulations for husbandry and handling of laboratory animals. Note that, compared to tail vein injections, retro-orbital injections require less technical expertise and may cause less distress in rodents^11^; however, tail vein injections appear to result in similar AAV distribution^17^.

CRITICAL When possible, verify viral transduction and transgene expression *in vitro* before systemic administration. Note that co-injecting AAVs with other substances (e.g., dyes) could affect infectivity *in vivo* and should be tested independently.

CRITICAL Re-titer viruses before injection if more than 1 month has passed since titration; this will ensure that animals are administered the most accurate dose possible.

1. Determine the dose of virus to administer per mouse (see Experimental design section for recommendations). Divide the dose (vg) by the titer (vg/ml) (**Supplementary Table 4**, cell G27) to calculate the volume of virus needed to inject one mouse. In a screw-cap vial, prepare a master mix of virus based on the number of animals to be injected; briefly vortex each virus and master mix 1-2 s before use. Transport the virus on ice once it is ready for injection. CAUTION Do not inject more than 10% of the mouse blood volume, which corresponds to 150 μl for a 25 g mouse. CRITICAL STEP Depending on the user, it is easiest to inject 40-80 μl/mouse. If less than 40 μl/mouse is required, use DPBS or saline to dilute the virus such that a larger volume is injected. If more than 80 μl/mouse is required, it may be more convenient to re-concentrate the virus or perform two separate injections; follow institutional guidelines for multiple eye injections. Virus will be lost in the event of an unsuccessful injection; therefore, prepare more master mix than is required. CRITICAL STEP To reduce the chance of contaminating the virus stock, avoid using the original virus stock; only bring an aliquot of what is needed for the injection(s). TROUBLESHOOTING
2. Assemble the anesthesia system53 inside the biosafety cabinet.
3. Remove the mouse from its cage and place it in the induction chamber. Anesthetize the mouse with 1-5% isoflurane. CAUTION Isoflurane must be handled according to federal, state, and local regulations.
4. While the mouse is being induced, load an insulin syringe with virus. Remove the dead space in the syringe barrel by gently ejecting the virus back into the tube such that air bubbles are expelled. Load the syringe again and repeat the procedure until no bubbles remain in the barrel. CAUTION Introducing air into the vascular system can be fatal. CRITICAL STEP Introducing air into the virus may cause protein denaturation; perform this step gently and only until no bubbles remain in the syringe barrel.
5. Remove the anesthetized mouse from the induction chamber. Place the animal in a prone position on a small stack of paper towels. Position the mouse such that its head is situated on the same side as the operator’s dominant hand. Place the nose cone on the mouse to maintain anesthesia.
6. Use the index finger and thumb on the non-dominant hand to draw back the skin above and below the eye, causing the eye to slightly protrude from the socket^11^. With the dominant hand, insert the needle, bevel down, at a 30-45° angle into the medial canthus and through the conjunctival membrane. The needle should be positioned behind the globe of the eye in the retro-orbital sinus. Slowly release the virus into the sinus and gently remove the needle. CAUTION Assess anesthetic depth by loss of pedal reflex (via toe pinch) before inserting the needle into the retro-orbital sinus. Any movement of the eye or skin when the needle is inserted indicates incorrect needle placement. Keep hands out of the path of the needle to prevent accidental exposure to AAVs. Do not recap needles; discard into an approved biohazardous sharps container immediately after use. CRITICAL STEP No liquid should leak out of the eye after viral delivery; likewise, little to no bleeding should be observed. TROUBLESHOOTING
7. Following viral injection, apply mild pressure to the eyelid. Apply 1-2 drops of proparacaine to the corneal surface to provide local analgesia. After recovery from anesthesia, place the mouse in a clean cage. CAUTION Monitor the eye daily after injection for 2 d, or according to institutional guidelines.

### Evaluation of transgene expression

TIMING Variable; see Experimental design section

CAUTION Follow appropriate institutional and governmental guidelines and regulations for husbandry and handling of laboratory animals.

1. Euthanize animals after sufficient time has passed for viral transduction and protein expression (see Experimental design section for recommendations). Assess fluorescent protein labeling (if applicable) in the tissue(s) of interest using standard histological methods for thin slices or tissue clearing (e.g., CLARITY^24^ or Sca/eSQ^25^) for thick tissue samples. Ensure that the chosen clearing protocol is compatible with the fluorescent protein(s) under investigation (see Experimental design and Anticipated Results sections for details). For experiments without fluorescent labels, evaluate transgene expression using molecular (e.g., qPCR or Western blot), histological (e.g. with antibodies, small molecule dyes, or molecular probes) or functional (e.g., optical imaging) methods relevant to the experimental aims. TROUBLESHOOTING

##### TIMING

Refer to **Figure 6a** for a timeline of the procedure.

Steps 1-3, Triple transient transfection of HEK293T cells: 1-2 h

Steps 4-14, AAV harvest: 5 d

Steps 15-31, AAV purification: 1 d

Steps 32-42, AAV titration: 1 d

Steps 43-49, Intravenous (retro-orbital) injection: 2-5 min per mouse, excluding set-up and clean-up time

Step 50, Evaluation of transgene expression: Variable

## ANTICIPATED RESULTS

### AAV production

For capsids that package well (i.e., AAV-PHP.B, AAV-PHP.eB, and AAV-PHP.S), the AAV production protocol typically yields over 1 × 10^12^ vg per 150 mm dish^2,3^ (**Supplementary Table 1**). Production efficiency can be determined for each virus in Step 42 (**Supplementary Table 4**, cell K27). Note that yields may vary from prep to prep and genome to genome. Users can gauge production efficiency for each experiment by running a positive control (e.g., pAAV-CAG-eYFP (Addgene ID 104055)).

### Evaluation of transgene expression

For most applications, users can expect to assess transduced cells beginning 2 or more weeks after intravenous injection (see Experimental design section for details). The chosen method for evaluating transgene expression will vary from user to user and may involve molecular, histological, and/or functional approaches (Step 50). We typically use fluorescent reporters to assess gene expression in thick (≥100 μm), cleared tissue samples; below, we discuss expected results for applications presented here (**Figs. 2-4**) and in our previous work^2,3^.

Commonly used reporters like GFP, eYFP, and tdTomato show strong fluorescent labeling in PACT- and PARS-cleared tissues, enabling whole-organ and thick-tissue imaging of transgene expression^3,22,24^. Most markers, including mTurquoise2, mNeonGreen, and mRuby2, can also be detected after mounting labeled tissues in optical clearing reagents like RIMS^24^ (**Fig. 3b**), Sca/eSQ^25^ (**Figs. 3a, 3c-d**, and **4b-d**), and Prolong Diamond Antifade (Thermo Fisher Scientific, cat. no. P36965)^2^. Depending on the rAAV genome, fluorescent proteins may be localized to distinct cellular compartments including the nucleus (via NLS) (**Fig. 2**), cytosol (**Figs. 3a-c** and **4b**), and cell membrane (via farnesylation^55^ or fusion to a membrane protein like ChR2) (**Figs. 3d** and **4d**). Some fluorescent proteins are susceptible to photobleaching. For example, mRuby2 may bleach over extended imaging sessions; tdTomato exhibits similar spectral properties and may be a more suitable alternative given its photostability^56^. Note that autofluorescent lipofuscin accumulates in aging postmitotic tissues (e.g., the brain and heart)^57^ and may interfere with examining transduced cells; either reduce autofluorescence using histological methods^24^ or, if possible, inject younger adults (≤8 weeks old) and determine the minimum time required for transgene expression.

Regardless of the approach used to evaluate gene expression, cell type-specific promoters should be verified at this stage in the protocol; we typically assess cell morphology and/or use antibody staining to confirm specificity (**Fig. 2b-c**).

### Conclusion

In summary, we present a comprehensive protocol for the production and administration of AAV-PHP viruses, which is also applicable to the production and systemic delivery of other AAV serotypes. We have validated the ability of AAV-PHP variants to provide efficient and noninvasive gene delivery to specific cell populations throughout the body. Together, this AAV toolbox equips users with the resources needed for a variety of applications across the biomedical sciences.

**Video 1. Steps 16 and 18 - Pouring the density gradient and loading the virus.**

**Video 2. Steps 26-27 - Collecting the virus.**

---

**Supplementary Table 1. AAV-PHP capsids for efficient transduction across specific organs and cell populations.** Species, organs, and cell populations examined to-date following intravenous administration of AAV-PHP viruses. To restrict gene expression to distinct cell types, use rAAV genomes with cell type-specific gene regulatory elements and/or Cre- or Flp-dependent recombination schemes (**Figs. 2-4**). Given the poor production efficiency of AAV-PHP.A, we suggest using AAV-PHP.eB to target astrocytes. Use an astrocyte promoter, such as GFAP, to drive transgene expression (**Fig. 2**).

**Supplementary Table 2. Transfection calculator**. This is an interactive calculator and provided as an Excel file (see Step 2 and REAGENT SETUP).

**Supplementary Table 3. Pouring the iodixanol density step solutions**. Determine the number of gradients needed and prepare the iodixanol density step solutions (see REAGENT SETUP for details).

**Supplementary Table 4. Titration calculator.** This is an interactive calculator and provided as an Excel file (see Step 42 and REAGENT SETUP).

## ACKNOWLEDGEMENTS

We thank Keith Beadle for contributions to the development of the protocol and for AAV production troubleshooting support. We also thank Mark Ladinsky for preparing transmission electron microscopy samples and for the image shown in **Figure 6**. This work was primarily supported by the National Institutes of Health (NIH) through grants to V.G.: Director’s New Innovator DP2NS087949 and PECASE; SPARC OT2OD023848-01 (to V.G. and S.K.); National Institute on Aging R01AG047664; BRAIN U01NS090577; and the Defense Advanced Research Projects Agency (DARPA) Biological Technologies Office (BTO; to V.G. and B.E.D.). Additional funding included the NSF NeuroNex Technology Hub 1707316 (to V.G.), the Curci Foundation (to V.G.), the Beckman Institute (to V.G. and B.E.D.), and the Rosen Center (to V.G.) at Caltech. V.G. is a Heritage Principal Investigator supported by the Heritage Medical Research Institute. R.C.C. was supported by an American Heart Association Postdoctoral Fellowship 17POST33410404. C.M.C was funded by the National Institute on Aging F32AG054101 and P.S.R. was funded by the National Heart, Lung, and Blood Institute F31HL127974.

## AUTHOR CONTRIBUTIONS

R.C.C. and V.G. wrote the manuscript with input from all coauthors. R.C.C., S.R.K., K.Y.C., C.M.C., M.J.J., P.S.R., J.D.T., S.K., B.E.D., and V.G. designed and performed experiments, analyzed the data, and prepared figures. V.G. supervised all aspects of the project. All authors edited and approved the manuscript.

## REFERENCES

1 Samulski, R. J. & Muzyczka, N. in Annual Review of Virology, Vol 1 Vol. 1 Annual Review of Virology (ed L. W. Enquist) 427–451 (2014).

2 Chan, K. Y. et al. Engineered AAVs for efficient noninvasive gene delivery to the central and peripheral nervous systems. Nature Neuroscience 20, 1172-+, doi:10.1038/nn.4593 (2017).

3 Deverman, B. E. et al. Cre-dependent selection yields AAV variants for widespread gene transfer to the adult brain. Nature Biotechnology 34, 204-+, doi:10.1038/nbt.3440 (2016).

4 Reed, S. E., Staley, E. M., Mayginnes, J. P., Pintel, D. J. & Tullis, G. E. Transfection of mammalian cells using linear polyethylenimine is a simple and effective means of producing recombinant adeno-associated virus vectors. Journal of Virological Methods 138, 85–98, doi:10.1016/j.jviromet.2006.07.024 (2006).

5 Wright, J. F. Transient Transfection Methods for Clinical Adeno-Associated Viral Vector Production. Human Gene Therapy 20, 698–706, doi:10.1089/hum.2009.064 (2009).

6 Xiao, X., Li, J. & Samulski, R. J. Production of high-titer recombinant adeno-associated virus vectors in the absence of helper adenovirus. Journal of Virology 72, 2224–2232 (1998).

7 Ayuso, E. et al. High AAV vector purity results in serotype- and tissue-independent enhancement of transduction efficiency. Gene Therapy 17, 503–510, doi:10.1038/gt.2009.157 (2010).

8 Grieger, J. C., Choi, V. W. & Samulski, R. J. Production and characterization of adeno-associated viral vectors. Nature Protocols 1, 1412–1428, doi:10.1038/nprot.2006.207 (2006).

9 Zolotukhin, S. et al. Recombinant adeno-associated virus purification using novel methods improves infectious titer and yield. Gene Therapy 6, 973–985, doi:10.1038/sj.gt.3300938 (1999).

10 Gray, S. J. et al. Production of recombinant adeno-associated viral vectors and use in in vitro and in vivo administration. Current protocols in neuroscience **Chapter 4**, Unit 4.17, doi:10.1002/0471142301.ns0417s57 (2011).

11 Yardeni, T., Eckhaus, M., Morris, H. D., Huizing, M. & Hoogstraten-Miller, S. Retro-orbital injections in mice. Lab Animal 40, 155–160 (2011).

12 Gray, S. J. et al. Preclinical Differences of Intravascular AAV9 Delivery to Neurons and Glia: A Comparative Study of Adult Mice and Nonhuman Primates. Molecular Therapy 19, 1058–1069, doi:10.1038/mt.2011.72 (2011).

13 Chakrabarty, P. et al. Capsid Serotype and Timing of Injection Determines AAV Transduction in the Neonatal Mice Brain. Plos One 8, doi:10.1371/journal.pone.0067680 (2013).

14 Maguire, C. A. et al. Mouse Gender Influences Brain Transduction by Intravascularly Administered AAV9. Molecular Therapy 21, 1469–1470 (2013).

15 Chen, Y. H., Chang, M. & Davidson, B. L. Molecular signatures of disease brain endothelia provide new sites for CNS-directed enzyme therapy. Nature Medicine 15, 1215–U1145, doi:10.1038/nm.2025 (2009).

16 Powell, S. K., Rivera-Soto, R. & Gray, S. J. Viral Expression Cassette Elements to Enhance Transgene Target Specificity and Expression in Gene Therapy. Discovery Medicine 19, 49–57 (2015).

17 Morabito, G. et al. AAV-PHP.B-Mediated Global-Scale Expression in the Mouse Nervous System Enables GBA1 Gene Therapy for Wide Protection from Synucleinopathy. Molecular Therapy, doi:http://dx.doi.org/10.1016/j.ymthe.2017.08.004 (2017).

18 Jackson, K. L., Dayton, R. D., Deverman, B. E. & Klein, R. L. Better Targeting, Better Efficiency for Wide-Scale Neuronal Transduction with the Synapsin Promoter and AAV-PHP.B. Frontiers in Molecular Neuroscience 9, doi:10.3389/fnmol.2016.00116 (2016).

19 Allen, W. E. et al. Global Representations of Goal-Directed Behavior in Distinct Cell Types of Mouse Neocortex. Neuron 94, 891-+, doi:10.1016/j.neuron.2017.04.017 (2017).

20 Hillier, D. et al. Causal evidence for retina-dependent and -independent visual motion computations in mouse cortex. Nature Neuroscience 20, 960-+, doi:10.1038/nn.4566 (2017).

21 Resendez, S. L. et al. Visualization of cortical, subcortical and deep brain neural circuit dynamics during naturalistic mammalian behavior with head-mounted microscopes and chronically implanted lenses. Nature Protocols 11, 566–597, doi:10.1038/nprot.2016.021 (2016).

22 Yang, B. et al. Single-Cell Phenotyping within Transparent Intact Tissue through Whole-Body Clearing. Cell 158, 945–958, doi:10.1016/j.cell.2014.07.017 (2014).

23 Richardson, D. S. & Lichtman, J. W. Clarifying Tissue Clearing. Cell 162, 246–257, doi:10.1016/j.cell.2015.06.067 (2015).

24 Treweek, J. B. et al. Whole-body tissue stabilization and selective extractions via tissue-hydrogel hybrids for high-resolution intact circuit mapping and phenotyping. Nature Protocols 10, 1860–1896, doi:10.1038/nprot.2015.122 (2015).

25 Hama, H. et al. Sca/eS: an optical clearing palette for biological imaging. Nature Neuroscience 18, 1518-+, doi:10.1038/nn.4107 (2015).

26 Choi, J. H. et al. Optimization of AAV expression cassettes to improve packaging capacity and transgene expression in neurons. Molecular Brain 7, doi:10.1186/1756-6606-7-17 (2014).

27 Paterna, J. C., Moccetti, T., Mura, A., Feldon, J. & Bueler, H. Influence of promoter and WHV post-transcriptional regulatory element on AAV-mediated transgene expression in the rat brain. Gene Therapy 7, 1304–1311, doi:10.1038/sj.gt.3301221 (2000).

28 Xu, R. et al. Quantitative comparison of expression with adeno-associated virus (AAV-2) brain-specific gene cassettes. Gene Therapy 8, 1323–1332, doi:10.1038/sj.gt.3301529 (2001).

29 de Leeuw, C. N. et al. rAAV-compatible MiniPromoters for restricted expression in the brain and eye. Molecular Brain 9, doi:10.1186/s13041-016-0232-4 (2016).

30 Gray, S. J. et al. Optimizing Promoters for Recombinant Adeno-Associated Virus-Mediated Gene Expression in the Peripheral and Central Nervous System Using Self-Complementary Vectors. Human Gene Therapy 22, 1143–1153, doi:10.1089/hum.2010.245 (2011).

31 Chamberlain, K., Riyad, J. M. & Weber, T. Expressing Transgenes That Exceed the Packaging Capacity of Adeno-Associated Virus Capsids. Human Gene Therapy Methods 27, 1–12, doi:10.1089/hgtb.2015.140 (2016).

32 Broderick, J. A. & Zamore, P. D. MicroRNA therapeutics. Gene Therapy 18, 1104–1110, doi:10.1038/gt.2011.50 (2011).

33 Xie, J. et al. MicroRNA-regulated, Systemically Delivered rAAV9: A Step Closer to CNS-restricted Transgene Expression. Molecular Therapy 19, 526–535, doi:10.1038/mt.2010.279 (2011).

34 Nayak, S. & Herzog, R. W. Progress and prospects: immune responses to viral vectors. Gene Therapy 17, 295–304, doi:10.1038/gt.2009.148 (2010).

35 Gao, K. et al. Empty virions in AAV8 vector preparations reduce transduction efficiency and may cause total viral particle dose-limiting side effects. Molecular Therapy-Methods & Clinical Development 1, doi:10.1038/mtm.2013.9 (2014).

36 Mingozzi, F. & High, K. A. Immune responses to AAV vectors: overcoming barriers to successful gene therapy. Blood 122, 23–36, doi:10.1182/blood-2013-01-306647 (2013).

37 Strobel, B., Miller, F. D., Rist, W. & Lamla, T. Comparative Analysis of Cesium Chloride- and Iodixanol-Based Purification of Recombinant Adeno-Associated Viral Vectors for Preclinical Applications. Human Gene Therapy Methods 26, 147–157, doi:10.1089/hgtb.2015.051 (2015).

38 Lerner, T. N. et al. Intact-Brain Analyses Reveal Distinct Information Carried by SNc Dopamine Subcircuits. Cell 162, 635–647, doi:10.1016/j.cell.2015.07.014 (2015).

39 Tervo, D. G. R. et al. A Designer AAV Variant Permits Efficient Retrograde Access to Projection Neurons. Neuron 92, 372–382, doi:10.1016/j.neuron.2016.09.021 (2016).

40 Cai, D., Cohen, K. B., Luo, T., Lichtman, J. W. & Sanes, J. R. Improved tools for the Brainbow toolbox. Nature Methods 10, 540-+, doi:10.1038/nmeth.2450 (2013).

41 Chang, R. B., Strochlic, D. E., Williams, E. K., Umans, B. D. & Liberles, S. D. Vagal Sensory Neuron Subtypes that Differentially Control Breathing. Cell 161, 622–633, doi:10.1016/j.cell.2015.03.022 (2015).

42 Williams, E. K. et al. Sensory Neurons that Detect Stretch and Nutrients in the Digestive System. Cell 166, 209–221, doi:10.1016/j.cell.2016.05.011 (2016).

43 Bruegmann, T. et al. Optogenetic control of heart muscle in vitro and in vivo. Nat Methods 7, 897–900, doi:10.1038/nmeth.1512 (2010).

44 Guettier, J. M. et al. A chemical-genetic approach to study G protein regulation of beta cell function in vivo. Proceedings of the National Academy of Sciences of the United States of America 106, 19197–19202, doi:10.1073/pnas.0906593106 (2009).

45 Jain, S. et al. Chronic activation of a designer G(q)-coupled receptor improves beta cell function. The Journal of clinical investigation 123, 1750–1762, doi:10.1172/jci66432 (2013).

46 Li, J. H. et al. A novel experimental strategy to assess the metabolic effects of selective activation of a G(q)-coupled receptor in hepatocytes in vivo. Endocrinology 154, 3539–3551, doi:10.1210/en.2012-2127 (2013).

47 Ran, F. A. et al. In vivo genome editing using Staphylococcus aureus Cas9. Nature 520, 186–191, doi:10.1038/nature14299 (2015).

48 Yang, Y. et al. A dual AAV system enables the Cas9-mediated correction of a metabolic liver disease in newborn mice. Nat Biotechnol 34, 334–338, doi:10.1038/nbt.3469 (2016).

49 Senis, E. et al. CRISPR/Cas9-mediated genome engineering: an adeno-associated viral (AAV) vector toolbox. Biotechnology journal 9, 1402–1412, doi:10.1002/biot.201400046 (2014).

50 Yang, Q. et al. AAV-based shRNA silencing of NF-kappaB ameliorates muscle pathologies in mdx mice. Gene Ther 19, 1196–1204, doi:10.1038/gt.2011.207 (2012).

51 Petri, K. et al. Comparative next-generation sequencing of adeno-associated virus inverted terminal repeats. Biotechniques 56, 269-+, doi:10.2144/000114170 (2014).

52 Masters, J. R. & Stacey, G. N. Changing medium and passaging cell lines. Nature Protocols 2, 2276–2284, doi:10.1038/nprot.2007.319 (2007).

53 Database, J. S. E. in Lab Animal Research (JoVE, Cambridge, MA, 2017).

54 Lock, M. et al. Rapid, Simple, and Versatile Manufacturing of Recombinant Adeno-Associated Viral Vectors at Scale. Human Gene Therapy 21, 1259–1271, doi:10.1089/hum.2010.055 (2010).

55 Hancock, J. F., Cadwallader, K., Paterson, H. & Marshall, C. J. A CAAX OR A CAAL MOTIF AND A 2ND SIGNAL ARE SUFFICIENT FOR PLASMA-MEMBRANE TARGETING OF RAS PROTEINS. Embo Journal 10, 4033–4039 (1991).

56 Day, R. N. & Davidson, M. W. The fluorescent protein palette: tools for cellular imaging. Chemical Society Reviews 38, 2887–2921, doi:10.1039/b901966a (2009).

57 Gray, D. A. & Woulfe, J. Lipofuscin and aging: a matter of toxic waste. Science of aging knowledge environment:SAGE KE 2005, re1, doi:10.1126/sageke.2005.5.re1 (2005).

58 Dana, H. et al. Sensitive red protein calcium indicators for imaging neural activity. eLife 5, doi:10.7554/eLife.12727 (2016).

59 Kim, J. et al. mGRASP enables mapping mammalian synaptic connectivity with light microscopy. Nature Methods 9, 96–U139, doi:10.1038/nmeth.1784 (2012).

60 Kim, E. J. & Sheng, M. PDZ domain proteins of synapses. Nat. Rev. Neurosci. 5, 771–781, doi:10.1038/nrn1517 (2004).

